# Immunogenic Cell Death: the Key to Unlocking the Potential for Combined Radiation and Immunotherapy

**DOI:** 10.1101/2025.02.14.638342

**Authors:** Somiya Rauf, Alexandra Smirnova, Andres Chang, Yuan Liu, Yi Jiang

**Author notes:** Corresponding author(s). E-mail(s); Contributing authors.

## Abstract

Immunogenic cell death (ICD) enhances anti-tumor immunity by releasing tumor-associated antigens and activating the anti-tumor immune system response. However, its potential remains understudied in combination therapies. Here, we develop a mathematical model to quantify the role of ICD in optimizing the efficacy of combined radiotherapy (RT) and macrophage-based immunotherapy. Using preclinical murine data targeting the SIRP*α*-CD47 checkpoint, we show that RT alone induces minimal ICD, whereas disrupting the SIRP*α*-CD47 axis significantly enhances both phagocytosis and systemic immune activation. Our model predicts an optimal RT dose (6–8 Gy) for maximizing ICD, a dose-dependent abscopal effect, and a hierarchy of treatment efficacy, with SIRP*α*-knockout macrophages exhibiting the strongest tumoricidal activity. These findings provide a quantitative framework for designing more effective combination therapies, leveraging ICD to enhance immune checkpoint inhibition and radiotherapy synergy.

## Introduction

Immunotherapy has become a major player in the clinical management of cancer over the past two decades [1]. Immunogenic cell death (ICD), a form of cell death that results from insufficient cellular adaptation to specific stressors, activates the immune system, initiates an inflammatory response and facilitates the recognition of dying cells by the immune system, is central in the development of novel anticancer treatments [2]. ICD involves changes in the cell surface as well as the release of soluble mediators, such as the externalization of calreticulin and the release of ATP and HMGB1 [3–5]. These signals add to other damage-associated molecular patterns (DAMPs) released by stressed or injured cells, operate on a series of receptors expressed by antigen presenting cells (APCs, such as dendritic cells and macrophages) to stimulate the presentation of tumor antigens to cytotoxic T cells [6], setting off anti-tumor immune response and additional ICD. This feedback loop of events not only eliminates the dying tumor cells, but also targets therapy-resistant cancer stem cells, making the induction of ICD a critical mechanism underlying the success of many cancer therapies [6, 7].

Moreover, the synergy between radiotherapy (RT) and immunotherapy has been shown to significantly enhance therapeutic results [8]. Studies have demonstrated that radiation-induced ICD is dose-dependent, with higher doses more effectively inducing immunogenic signals [9]. However, most RT modalities, even when combined with immunotherapy (e.g., checkpoint inhibitors to augment antitumor immunity), cannot induce antitumor efficacy, especially when tumors are large, poorly immunogenic, and/or harbor a strong immunosuppressive milieu [10, 11].

Signal-regulatory protein *α* (SIRP*α*) is a key checkpoint that regulates macrophage activity and plays a pivotal role in phagocytosis [12]. Its interaction with CD47–a transmembrane protein over-expressed in cancer cells [13]– confers a “don’t eat me” signal, enables the cancer cells to evade immune surveillance [14].

Macrophage exhibits phenotypic plasticity, transitioning between M1 (classically activated, pro-inflammatory, and tumoricidal) and M2 (alternatively activated, anti-inflammatory, and tumor-promoting) macrophages [15–17]. The balance between M1 and M2 or M2-like phenotype for macrophages in the tumor microenvironment (TME) significantly influences cancer progression [18]. Efforts in blocking the SIRP*α*-CD47 interactions using anti-CD47 antibodies [19–21], anti-SIRP*α* antibodies [22], or SIRP*α*-Fc fusion proteins [23], have been shown to increase the M1/M2 ratio and enhance tumor eradication. Furthermore, SIRP*α*-knockout (KO) macrophages demonstrate enhanced phagocytic activities and tumor-targeting capabilities in pre-clinical models [24, 25], characterized by high inducible nitric oxide synthase (iNOS) expression, decreased CD206 level, and enhanced phatocytosis and antigen pre-sentation capacity [25]. These findings suggest that SIRP*α* deficiency promotes a pro-inflammatory and tumoricidal M1 phenotype, enhancing anti-tumor immunity.

Despite promising preclinical successes [25–28], translating SIRP*α*-CD47 blockade therapies into clinical applications remains challenging. A key knowledge gap exists in understanding how these immunotherapies interact with radiation therapy (RT), particularly macrophage-based approaches. Combining anti-SIRP*α* treatment with RT results in superior antitumor responses beyond either therapy alone by transforming the TME into a pro-inflammatory, tumoricidal niche [28]. Similar effects, including both local control and abscopal responses, have been observed in small cell lung cancer models following anti-CD47 treatment [26]. Moreover, RT in SIRP*α*-deficient mice induces robust pro-inflammatory responses , cultivating a tumoricidal microenvironment characterized by cytotoxic T cell, natural killer cell, and inflammatory neutrophil infiltration, while minimizing immunosuppressive factors [25].

Mathematical modeling of cancer immunotherapy has emerged as a critical approach for unraveling the complex interactions between tumor cells and the immune system. These models considered tumor cells, effector cells, cytokines, macrophages, and other immune cells [29–31]. Some consider the role of tumor-associated macrophages in tumor growth [32], aiming to predict drug delivery, radiation dose, and patient response. Prior models of macrophages distinguished M1 and M2 phenotypes and evaluated their contributions to controlling tumor growth and immunosuppression [33–35], suggesting Tumor associated macrophages (TAMs)-induced immunosuppression negatively impacts treatment efficacy. However, these models have not explored macrophages in the context of immune checkpoint inhibition via the SIRP*α*-CD47 checkpoint, or their combination with RT.

A great number of factors must coordinate for cancer immunotherapy to be effective. No single mathematical model has yet incorporated every aspect of every process involved in tumor growth *in vivo*, TME, immunotherapy, and RT. However, this level of complexity is not necessary for a model to be useful or predictive. In this study, we develop a minimal mathematical model that integrates tumor-immune interactions, RT, immunotherapies targeting the SIRP*α*-CD47 binding, and ICD. The model is calibrated using tumor growth data from wild-type (WT) and SIRP*α*^−*/*−^ mice with or without RT [25]. Our results indicate RT-induced ICD is minimal in WT mice but varies in SIRP*α*-deficient mice depending on radiation dose and tumor size. Furthermore, WT macrophages exhibit limited phagocytic activity, whereas disrupted SIRP*α*-CD47 binding enhances phagocytosis to different degrees. The calibrated model predicts the treatment effects using macrophages with SIRP*α*-KO, anti-SIRP*α*, CD47-KO, and anti-CD47, and quantitatively compare their treatment efficacy. Furthermore, it provides a comprehensive framework to assess treatment outcomes based on RT dose, ICD, and macrophage phagocytosis rate for all perturbations on the SIRP*α*-CD47 axis. Ultimately, our finding reveals that ICD plays a central role in effective cancer treatment through combined immunotherapy and RT.

## Results

### A new mathematical model for *in vivo* tumor growth under RT and macrophage-based immunotherapy

Our model considers three populations (cancer cells, effector cells, and macrophages) and four ways of cancer cell death (effector killing, phagocytosis by macrophage, radiation cell death, and immunogenic cell death), as illustrated in Figure 1. Radiation damaged cells activate the ICD pathway by activating the effector cells that kill more cancer cells (blue arrow in Figure 1A). Macrophages with disrupted CD47/SIRP*α* binding promote a highly pro-inflammatory TME and enhance ICD (orange arrows). Briefly, tumor growth follows a logistic dynamics and is either inhibited by effector cells or phagocytic macrophages. Effector cells infiltrate the tumor, become exhausted with reduced polyfunctionality, or cleared at a rate that we assume to be constant. Similarly, macrophages infiltrate, become exhausted from phagocytosing tumor cells or cleared at a rate assumed to be constant. RT reduces tumor cell survival in a dose-dependent manner, with immunogenic cell death proportional to the RT-induced cell death (Figure 1B).

**Figure 1.**
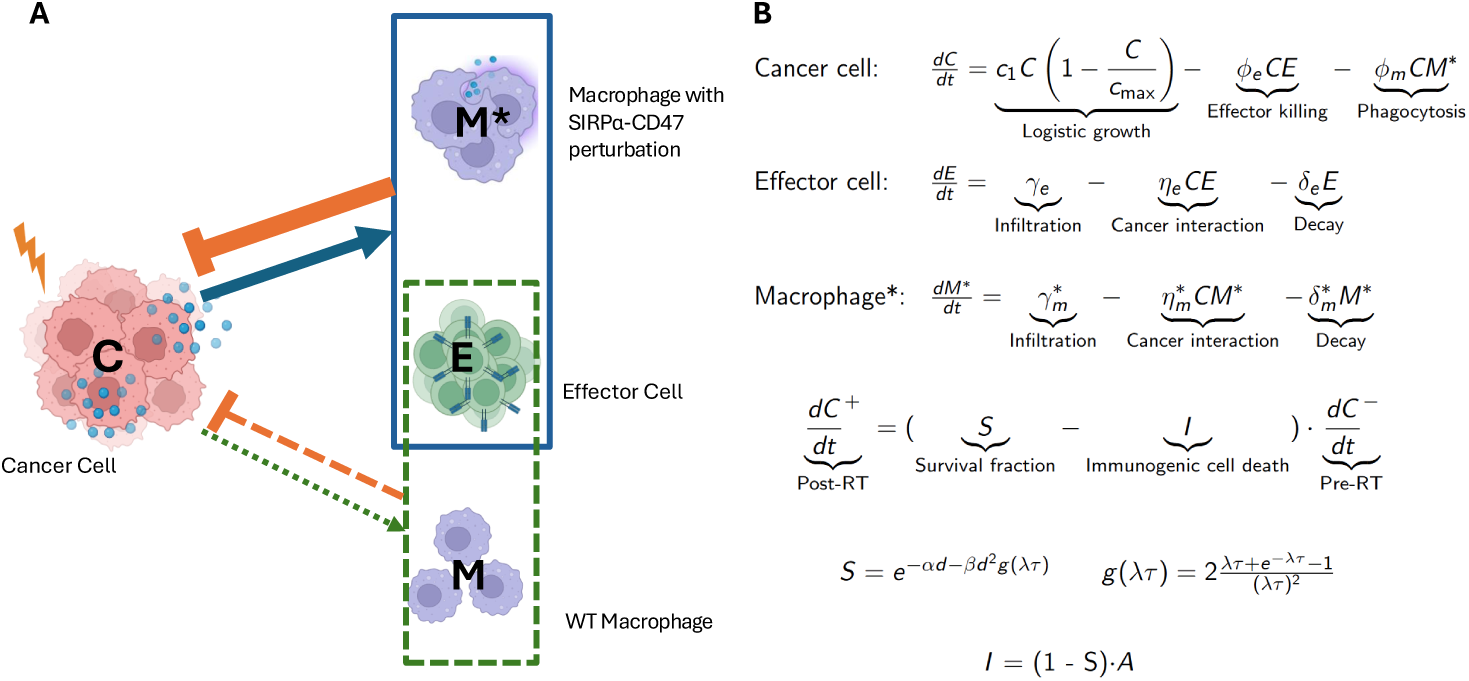
Overview of the mathematical model: **A.** Radiation induces minimal activation of the immunogenic cell death (ICD) pathway (dashed green arrow) with the WT macrophages. Macrophages with disrupted SIRP*α*-CD47 checkpoint respond to RT-induced DAMPs, activating ICD (solid blue arrow) that leads to enhanced tumor elimination (solid orange hammer-head). **B.** The model equations describe the dynamics of cancer cells (*C*), effector cells (*E*), and macrophages (*M*). Cancer cells after radiation follow a linear-quadratic (LQ) dose-dependent survival rate (*S*) and ICD *I* that is proportional to radiation damage (1 *− S*) and ICD activation rate *A*. Details are in the Methods and Supporting Information.

We calibrate the model using tumor growth data from a preclinical study using treatment-resistant colorectal adenocarcinoma (MC38) in WT (C57BL/6) and SIRP*α*^−*/*−^ mice, with and without local irradiation. Unless otherwise noted, all experimental data used in this paper were previously published [25]. This calibration is done in four steps of increasing complexity. We first find the parameters of tumor growth in WT mice (Table 1). We fix these parameters when we calibrate RT-related parameters of tumor growth in WT with RT (Supplementary Table 5, and SIRP*α*^−^ macrophage related parameters using tumor growth data in SIRP*α*^−*/*−^ mice without RT . We then calibrate for the last parameters for tumor growth in SIRP*α*^−*/*−^ mice with RT. The goodness of fit is evaluated using an error rate of *|fit − data|/data*. The calibration processing constraints possible explanations of experimental data, and the calibrated model allows for predictions. The prediction results reported below are calculated using the calibrated model without any free parameters. An important consideration is that *in vivo* tumor growth data, measured in volume, includes both cancer cells and infiltrated immune cells.

**Table 1.**
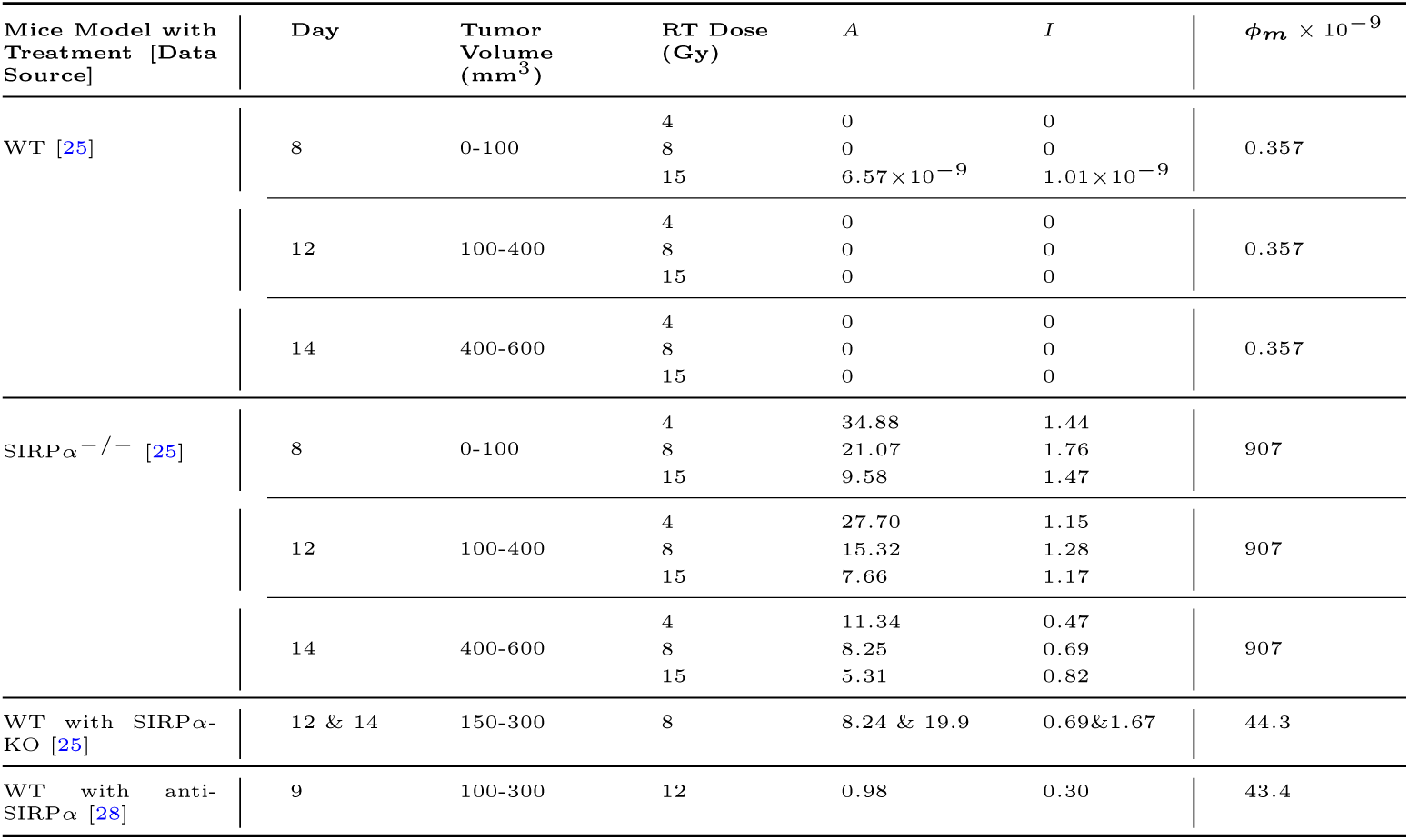
ICD and phagocytosis values for MC38 tumors across different SIRP*α*-CD47 treatments.

#### RT in WT mice invokes limited ICD

The model effectively replicates tumor growth data in WT mice with RT of various doses, where MC38 cells (5 *×* 10^5^) were subcutaneously engrafted into the right flank of WT mice and irradiation was delivered on day 8, 12, and 14, corresponding to tumor volumes of 0 *−* 100*mm*^3^ (small), 100 *−* 400*mm*^3^ (medium), and 400 *−* 600*mm*^3^ (large), respectively (data from [25], Figure 2A,B and C). RT delivered on later days and larger tumors results in less tumor reduction. For these tumor sizes and RT doses, immunogenic cell death values (*I* = (1*−S*)*A*) remain zero, except for small tumors receiving the largest dose (15 Gy), where *I ≈* 10^−9^ – negligible compared to ICD values when the SIRP*α*-CD47 pathway is disrupted (Table 1). These results suggest that in WT mice treated with RT, the impact of ICD is minimal, even for small tumors subjected to high-dose irradiation.

**Figure 2.**
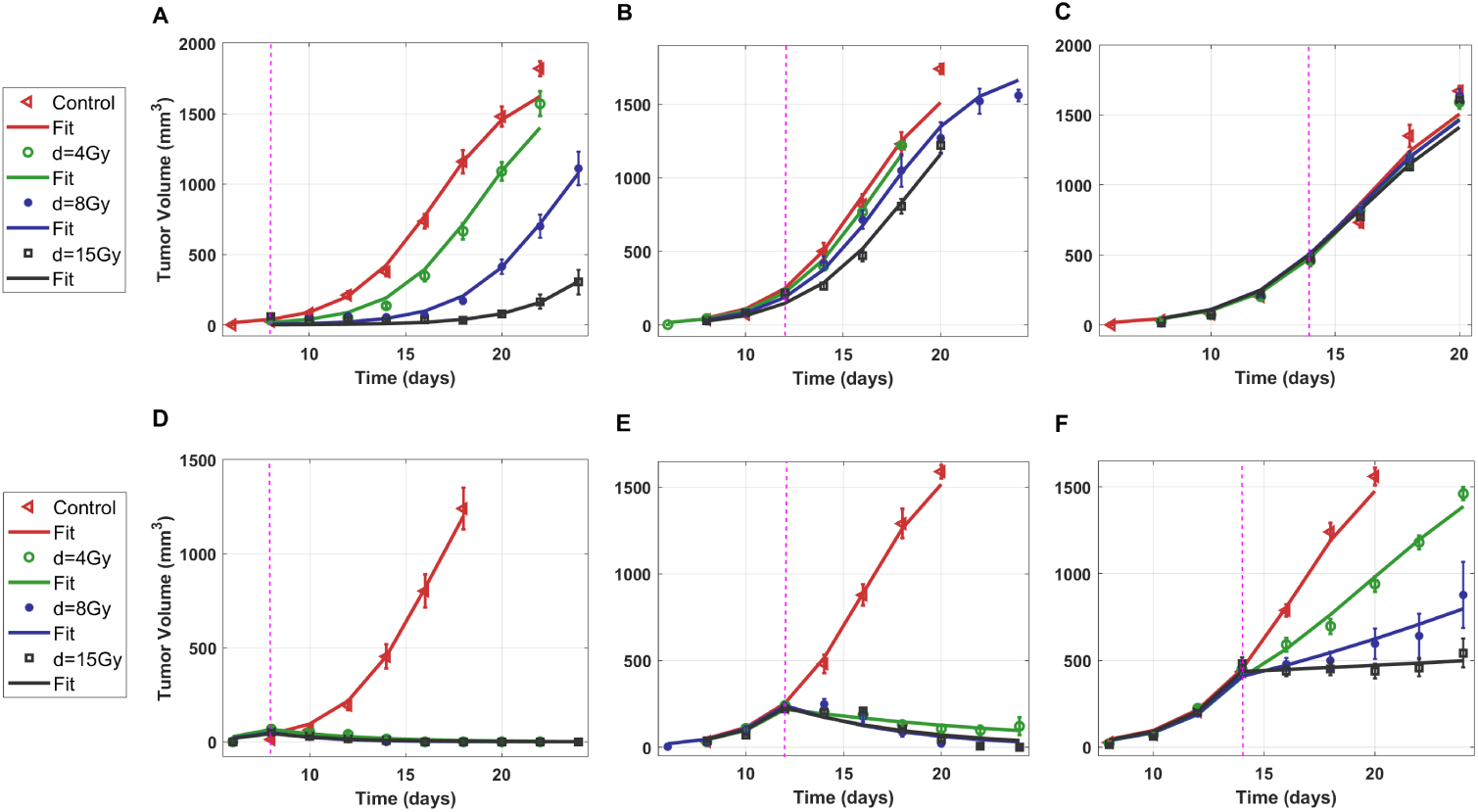
RT is ineffective for large tumors in WT mice but significantly reduces tumor growth in SIRP*α^−/−^* mice: **A-C.** In WT mice, small tumors (0 − 100*mm*^3^) respond well to higher doses of RT (**A**), while medium-sized tumors (100 − 400*mm*^3^) exhibit only a modest reduction even with large doses (B). Large tumors (400 − 600*mm*^3^) show minimal size eduction in size. **D-F.** In SIRP*α^−/−^* mice, RT effectively eliminates small tumors (**E**), remains effective for medium-sized tumors , **E**, and offers significant reduction for large tumors (**F**) depending on radiation dose.

#### ICD in SIRP*α*^−*/*−^ mouse depends on tumor size and radiation dose

In SIRP*α*^−*/*−^ mice, significant tumor reduction is evident following RT, even in large tumors (Figure 2D, E and F). This increased tumor reduction can be attributed to ICD, which is influenced by the interplay between RT cancer cell damage and effector cell activation. The corresponding ICD values for small, medium, and large tumors at RT dose of 4, 8, and 15 Gy are summarized in Table 1.

While the values for ICD activation *A* and ICD *I* vary across different tumor volumes and RT doses (Table 1), an interesting trend emerges: ICD consistently peaks at an RT dose of 8 Gy for all tumor sizes. This observation can be explained by the fundamental mechanism of ICD. Radiation-induced cancer cell damage (1 *− S*) increases with dose; however, ICD activation, *A*, decrease at higher doses (Table 1) due to RT-induced damage to immune cells. The interplay between these two opposing effects results in a biphasic dependence of ICD on radiation dose. Notably, previous studies have also showed that intermediate radiation doses provide optimal treatment effects in mice [36].

We explore the ICD (*I*) values across different tumor volumes and radiation doses to determine the effectiveness of RT in inducing ICD (Figure 3). This result is calculated by the spline interpolation of the immune activation parameter *A* in Table 1. Notably, in small and medium sized tumors, a progressive increase in immune activation is observed with increasing doses up to a threshold of 6 Gy. This observation suggests that optimal RT dose is close to 6 Gy for smaller tumors (*<* 400*mm*^3^). For large tumors, despite the increasing radiation doses, the rate of immune activation increase is more gradual till 10 Gy, suggesting that larger tumors may pose challenges in eliciting a strong immune-mediated anti-tumor effect through radiation alone. This underscores the importance of considering tumor burden when designing radiation doses for optimal ICD and highlights the potential need for combined therapies to overcome immune resistance in larger tumors.

**Figure 3.**
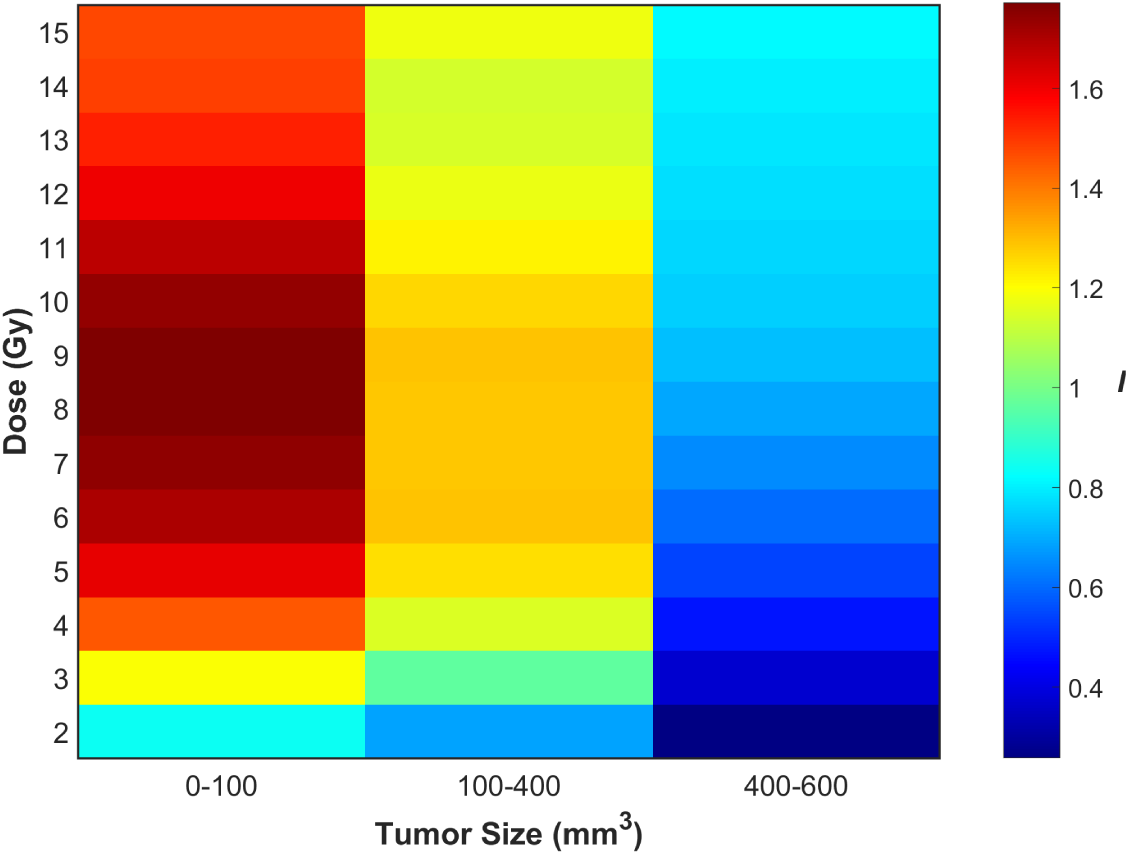
Optimizing RT doses to enhance ICD in SIRP*α^−/−^* mice: Immunogenic death induced (*I*) by RT in the SIRP*α*-/- mice model. The heat-map highlights high immune activation for intermediate doses for small and intermediate tumor volumes. For Large tumor the medium and high doses of RT are giving better results.

The sensitivity analysis of LQ model parameters in WT and SIRP*α*^−*/*−^ mice, focusing on different tumor sizes, reveals that these RT parameters have minimal impact on tumor response in WT mice, while a more pronounced effect is observed in SIRP*α*^−*/*−^ mice (Supplementary Figure 4).

#### Combined with RT, SIRP*α*-deficient macrophages effectively reduce tumor growth via ICD

In SIRP*α*^−*/*−^ mice, tumor responsiveness to RT is contingent upon the presence of SIRP*α*^−^ macrophages [25]. Depletion of intratumoral (i.t.) macrophages using either Cl2MDA liposomes or an antibody against the CSF1 receptor (*α*CSF1R) abolishes the efficacy of RT in SIRP*α*^−*/*−^ mice [25]. Our model accurately predicted this effect without any fitting parameters (Figure 4A.)

**Figure 4.**
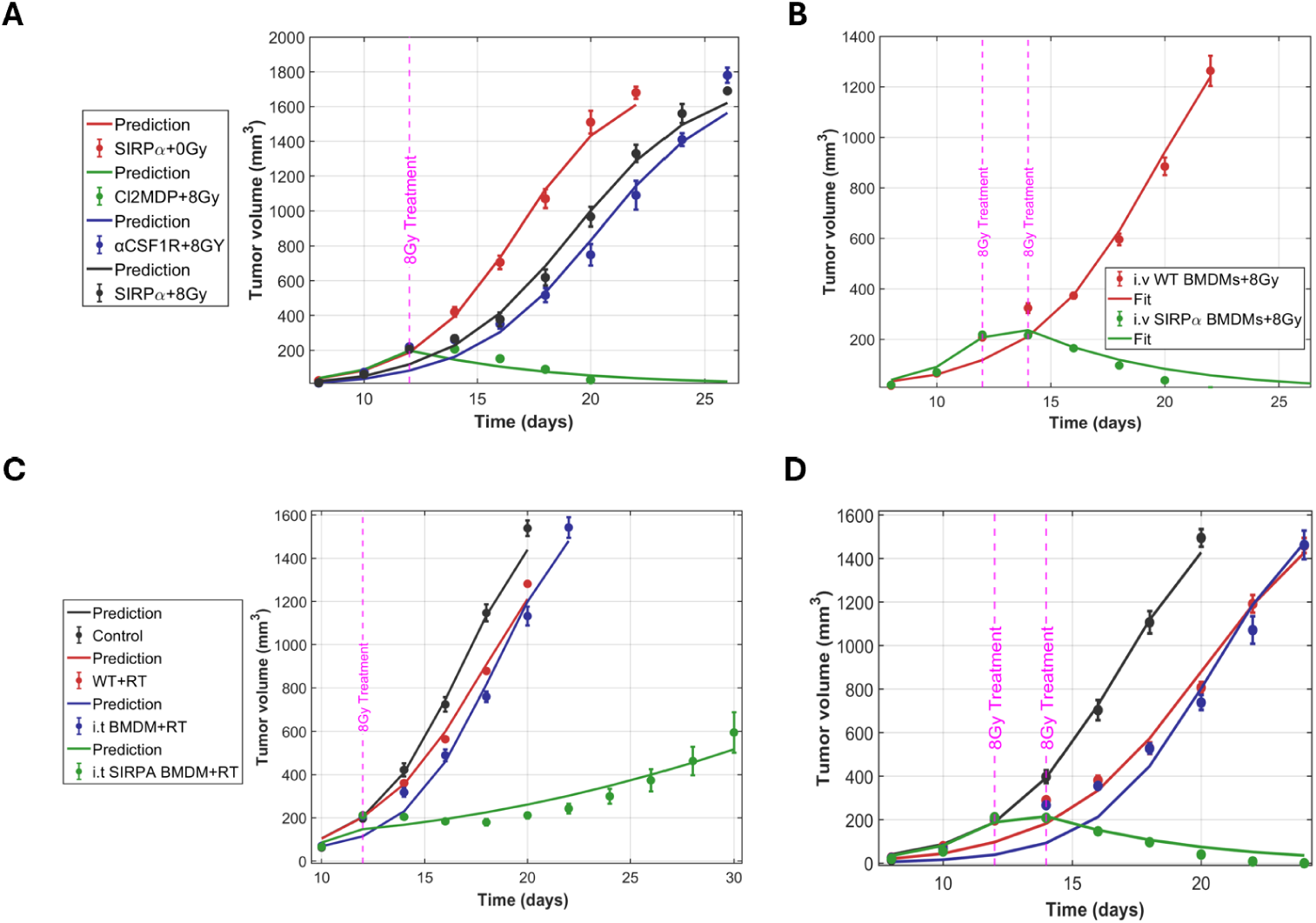
SIRP*α*-deficient macrophages reduce tumor growth in WT mice via ICD. **A.** Mathematical model predicts no tumor killing in SIRP*α^−/−^* mice with RT when macrophages are depleted. **B.** The mathematical model captures tumor elimination by twice i.v. injection of SIRP*α*-deficient macrophages with RT of 8 Gy. **C, D.** The model predicts that single and dual intratumoral (i.t.) treatments with BMDM and SIRP*α^−/−^* macrophages in WT mice with 8 Gy irradiation yield comparable tumor elimination as in i.v. injection case.

SIRP*α*-deficient macrophages can have complete anti-tumor treatment effects in WT mice when injected intravenously (i.v.) accompanied by RT [25](Figure 4B). We allow only two free parameters for fitting these experimental data: immune activation *A* and marcophage phagocytosis rate *ϕ_m_*. We find that for the first i.v. injection on day 12 with 8 Gy RT, *A*_1_ = 8.247 (corresponding to *I* = 0.69), and the second i.v. injection with RT on day 14, *A*_2_ = 19.95 (*I* = 1.67). The immune activation for the second treatment is much higher than the first, suggesting immune priming is at work. Macrophage phagocytosis rate *ϕ_m_* = 4.43*×*10^−8^ remains the same for both treatments.

Next, using *A*_1_, *A*_2_, and *ϕ_m_* values from i.v. injection experiments, we predict the tumor response when WT mice receive intratumoral (i.t.) injection of WT (SIRP*α*^+^ bone marrow-derived macrophages (BMDM), and SIRP*α*^−^ macrophages, in conjunction with RT at 8 Gy [25]. The model successfully predicts the reduction in tumor growth with a single i.t. injection using *A*_1_ (Figure 4C), and twice i.t. injections using both *A*_1_ and *A*_2_ (Figure 4D). This confirms that immune activation for i.t. and i.v. injections results in the same values of ICD. Injecting SIRP*α*^−*/*−^ macrophages to WT mice transforms the TME into a state resembling that in SIRP*α*^−*/*−^ mice.

#### Model predicts phagocytosis order in SIRP*α*-CD47 checkpoint inhibitions

In both WT and SIRP*α*^−*/*−^ mice, the rate of tumor growth depends on the number of engrafted MC38 cells: a higher number of cells leads to more rapid tumor growth [25]. In WT mice (Figure 5A), We find that a smaller initial tumor size leads to larger intratumoral effector cell population, which in turn reduces tumor volume (Supplementary Figures 1C and 2A, C). The effector cells are responsible for lowering the tumor volume because macrophage phagocytosis rate is one order of magnitude lower than the reported M1 macrophage phagocytosis rate [37] (Supplementary Table 1). In addition, we show that tumor growth in WT mice is insensitive to macrophage-related parameters, suggesting that WT macrophages have little impact on the tumor volume change (Supplementary information and Supplementary Table 1). Moreover, while the growth rates of tumors are similar between WT and SIRP*α*^−*/*−^ mice when larger numbers of MC38 cells are engrafted, SIRP*α*-/- mice exhibits a significantly slower tumor growth rate with a smaller engraftment at 5 *×* 10^3^ cells/mouse (Figure 5B). Our model demonstrates that the phagocytosis rate of SIRP*α*^−^ macrophages is the primary factor accounting for this variance. Interestingly, this phagocytosis rate does not reduce tumor growth for large engraftments, due to a rapid decrease of macrophages in SIRP*α*^−*/*−^ mice (Supplementary Figure 2E).

**Figure 5.**
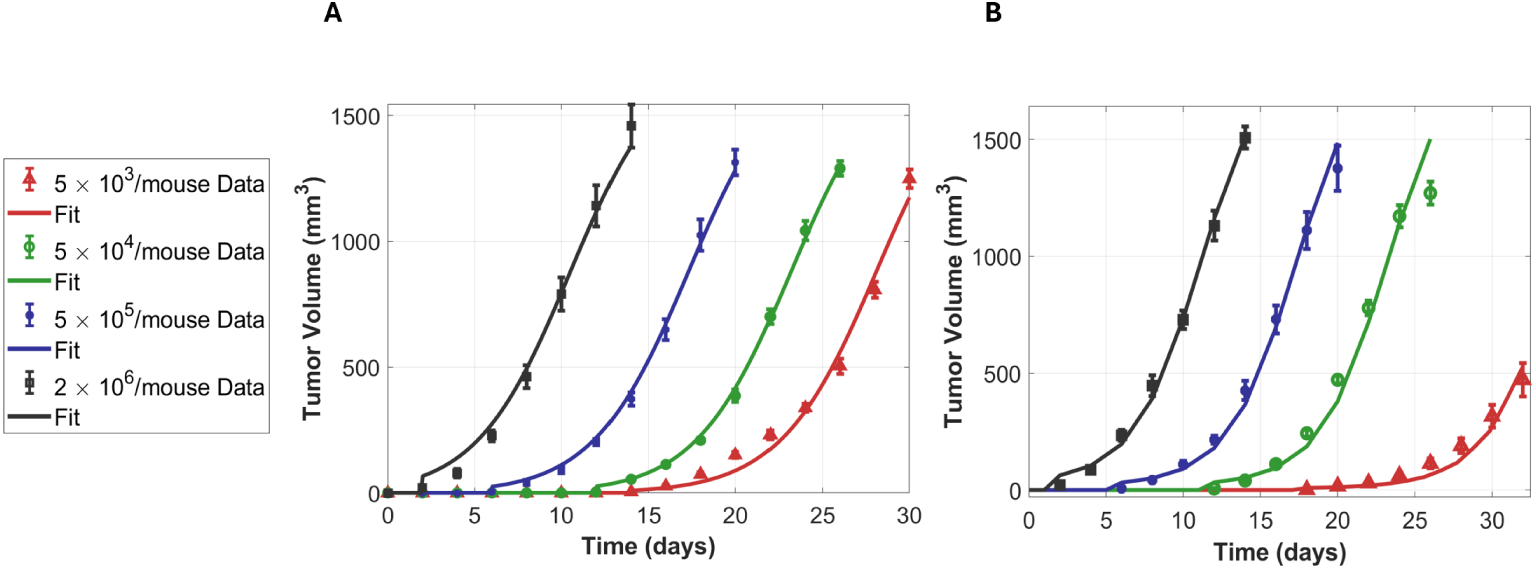
SIRP*α^−^*macrophages show higher phagocytosis rates than WT: Tumor growth patterns across four tumor cell engraftments in WT (A) and SIRP*α^−/−^* (B) mice, with model fitting (solid lines) and experimental data (dots with error bars) experimental data from [25].

Both anti-CD47 and CD47-KO promote phagocytosis by disabling the “don’t eat me” signal [38]. We find that inhibitions of SIRP*α*-CD47 checkpoint together with RT in WT mice increase the phagocytosis rate by two orders of magnitudes (Table 1). This observation aligns with experimental findings [25, 39], which report the enhanced phagocytic capability of SIRP*α*-deficient macrophages. Interestingly, macrophages in two different anti-SIRP*α* studies [27, 28], where RT of different doses were applied to tumors of different sizes, show the same phagocytosis rates. This suggest that checkpoint inhibition determined phagocytosis rate is independent of RT dose or tumor size. These results also allow us to rank the phagocytosis rates of macrophages with SIRP*α*-CD47 checkpoint inhibition, in descending order: SIRP*α*-KO, anti-SIRP*α*, and anti-CD47. We do not have MC38 tumor growth data with combined CD47-KO and RT in WT mice (Tables 1 and 2).

**Table 2.**
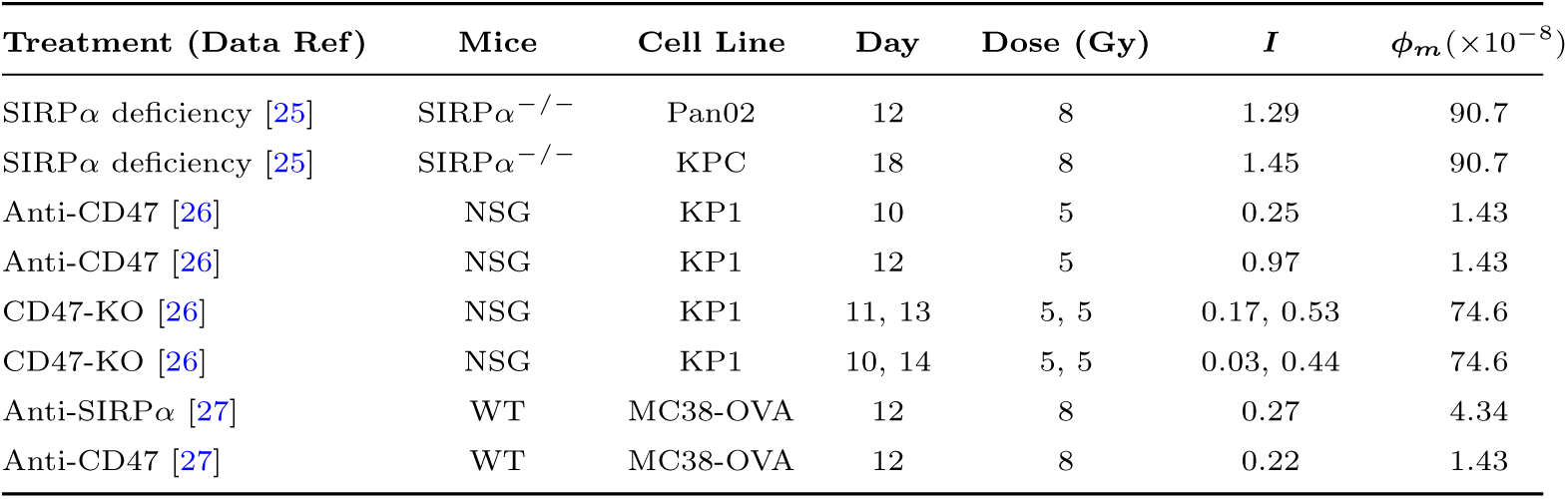
Summary of treatments in other mice models and other cell lines: Among the tested cell lines, the KPC cell line exhibited the strongest ICD response, surpassing MC38 and Pan02. Anti-CD47 therapy induced a high ICD compared to CD47-KO.

#### Model explains enhanced tumor clearance with combined RT and SIRP*α*-CD47 checkpoint inhibitions in other mouse and tumor models

To determine if our model is useful for only MC38 tumors in WT and SIRP*α*^−*/*−^ mice, we apply the model to other tumor and mouse models with combined RT and SIRP*α*-CD47 checkpoint inhibitions. We use tumor growth data for KP1 small cell lung cancer (SCLC) allografts in immunodeficient NSG mice under RT in combination with CD47 blockade (Figure 6A, B), data from [26], MC38-Ovalbumin (MC38-OVA) tumor in WT mice with RT and anti-CD47 or anti-SIRP*α* treatment (Figure 6C), data from[27], and MC38 tumor in WT mice with anti-SIRP*α* and high-dose radiotherapy (HRT) (Figure 6D) experimental data from [28]. In addition, we use growth data from pancreatic cancer cell lines Pan01 and KPC in WT and SIRP*α*^−*/*−^ mice (Figure 6E, F) [25]. Our model is able to fit for all tumor growth curves, showcasing its general applicability (Figure 6). Their phagocytosis rate (*ϕ_m_*) and ICD values (*I*) are summarized in Table 2. All other parameter values for different cell lines and mouse models are summarized in Supplementary Table 6 and 7.

**Figure 6.**
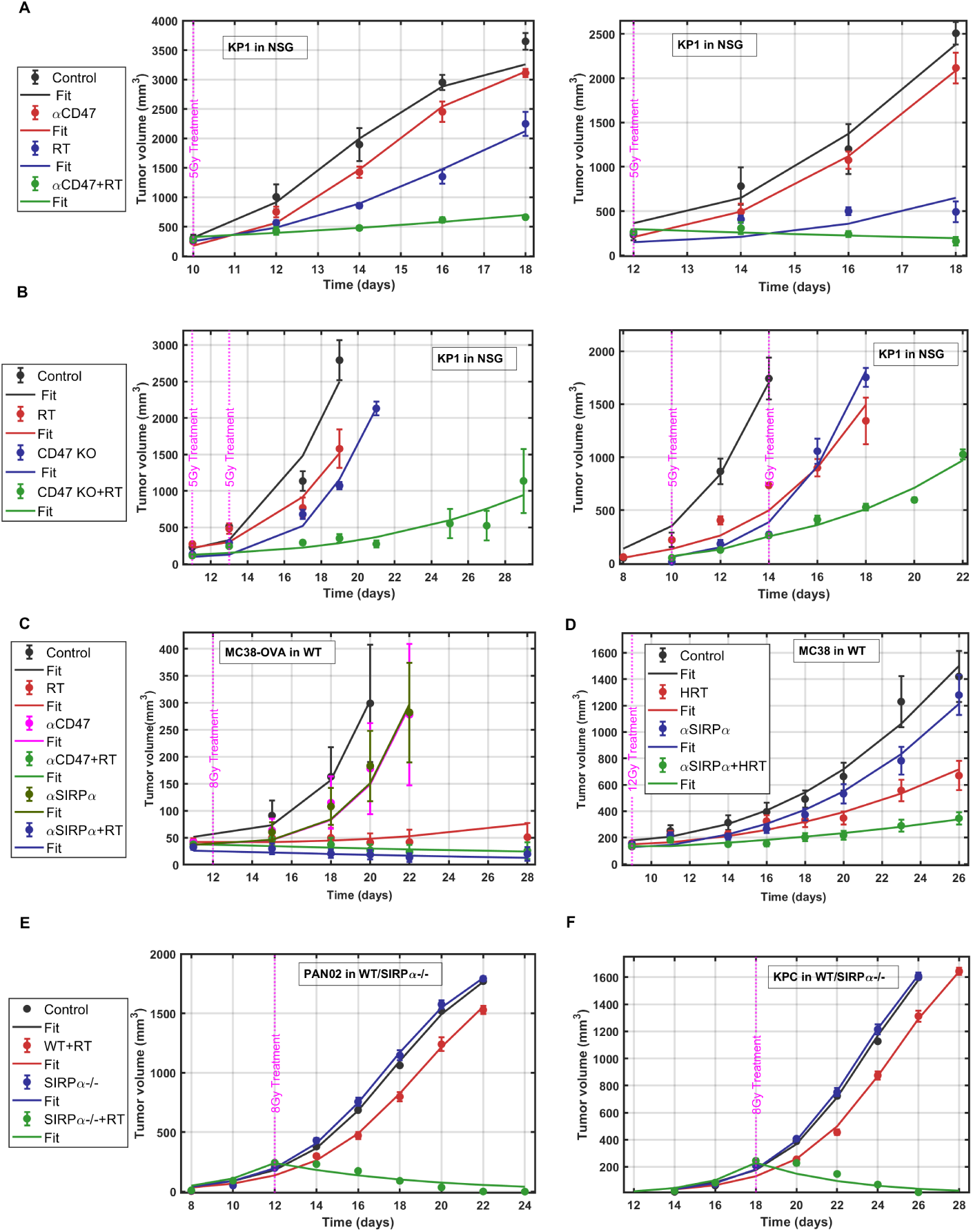
Model explains enhanced tumor clearance with combined SIRP*α*-CD47 check-point inhibitions and RT across various mouse and tumor models: **A.** KP1 tumors in NSG mice treated with anti-CD47 and RT, based on experimental data from [26]. **B.** KP1 tumors in NSG mice treated with CD47-KO with multiple RT, based on experimental data from [26]. **C.** MC38-OVA tumors in WT mice treated with anti-SIRP*α* and anti-CD47 and RT, based on experimental data from [27]. **D.** MC38 tumors in WT mice treated with anti-SIRP*α* and RT, based on experimental data from [28]. **E.** Pan02 tumor growth in WT and SIRP*α^−/−^* mice with RT, based on experimental data from [25]. **F.** KPC tumor growth in WT and SIRP*α^−/−^* mice with RT, based on experimental data from [25]. Model fitted parameters allow us to compare the macrophage phagocytosis rates and ICD values.

Because of substantial differences in genetic background, immune competency, and tumor microenvironment among various mouse models, direct comparisons of tumor growth across these models are confounded by multiple uncontrolled variables. Attempting to draw meaningful parallels or establish universal growth parameters from such heterogeneous systems often lacks scientific rigor and can lead to misleading conclusions. We therefore only compare within the same tumor cell lines or within the same mouse models.

The ICD value *I* in MC38 tumors in WT mice with combined RT and anti-CD47 treatment is significantly lower than that in SIRP*α*^−*/*−^ mice (Table 2). In addition, the ICD values for MC38 tumors across different SIRP*α*-CD47 checkpoint inhibitions in combination with RT in WT mice shows a clear pattern: earlier treatment leads to enhanced ICD (Table 1). Treatment of RT with SIRP*α*-KO achieves the highest ICD, followed in descending order by anti-SIRP*α*, anti-CD47, CD47-KO treatments, and control (no treatment). A second treatment elicits much higher ICD, suggesting that the immune system could be primed by the first treatment.

In SIRP*α*^−*/*−^ mice, immune activation due to the same irradiation dose (8 Gy) is higher for KPC tumor than Pan02 and MC38 tumors, suggesting that KPC cells can be more susceptible to SIRP*α*-based treatment under RT (Table 2).

For KP1 tumors in NSG mice, we find ICD decreasing with tumor size in combined anti-CD47 and RT treatment (Table 2), consistent with the trend in MC38 tumors in WT mice (Table 1).

Another recent study identified that irradiated colorectal cancer (CRC) cells use the ataxia-telangiectasia and Rad3-related (ATR)-mediated DNA double-strand break repair pathway to upregulate CD47 and PD-L1, which interacted with SIRP*α* and PD-1, respectively, to suppress phagocytosis and tumor-associated antigens (TAA) cross-presentation by APCs [27]. RT combined with anti-CD47/anti-SIRP*α* antibodies enhanced dendritic cell function and CD8 T-cell priming, and propagated the local tumoricidal activity of RT into vigorous systemic antitumor immunity [27]. Our model is able to fit their MC38-OVA tumor growth data in WT mice with RT and anti-CD47 or anti-SIRP*α* (Figure 6C) (Supplementary Table 7.) We find immune activation in case of anti-CD47 treatment (*A* = 0.27) is lower than the case of anti-SIRP*α* treatment (*A* = 0.33), suggesting that anti-SIRP*α* is more effective than anti-CD47 for MC38-OVA.

Our mathematical model also recapitulates MC38 tumor growth data in WT (C57BL/6) mice with anti-SIRP*α* and high-dose radiotherapy (HRT) of 12 Gy[28] (Figure 6D). Comparing the ICD activation *A* values (Supplementary Table 7), we observe that immunogenic activation with HRT is much lower in WT mice with anti-SIRP*α* treatment than in SIRP*α*-deficient mice.

For pancreatic cancer (Pan02 and KPC) growth data in WT and SIRP*α*^−*/*−^ mice, we fit *c*_1_, *c_max_* and *A*, (Figures 6E &F). We keep radio-sensitivity parameter same as MC38 cells to avoid parameter overfitting. Note that RT of 8 Gy are administered on day 12 for Pan02 and day 18 for KPC, respectively, to keep the same tumor sizes of 200*mm*^3^. We find *A* = 15.33 for Pan02, comparable similar to that of MC38 tumors, and *A* = 17.24 for KPC, results in higher ICD for KPC in comparison to MC38 and Pan02.

#### Model predicts abscopal effect in SIRP*α*^−*/*−^ **mice.**

The abscopal effect, although rare, is a systemic immune response that reduces tumor burden outside the irradiated area. Our mathematical model successfully predicts the abscopal effect in SIRP*α*^−*/*−^ mice and reproduces the tumor growth data reported in [25] without parameter fitting. Interestingly, the model demonstrates a complete reduction of abscopal tumors smaller than *<* 100*mm*^3^ (Figure 7A). However, for abscopal tumors larger than 200*mm*^3^, the reduction is less pronounced (Figure 7B), with some tumor cells surviving and leading to relapse one week post-treatment. Our model does not capture this relapse effectively, likely due to the simplified assumption of constant ICD values. A more sophisticated model incorporating adaptive immune dynamics may be needed to better describe abscopal tumor relapse.

**Figure 7.**
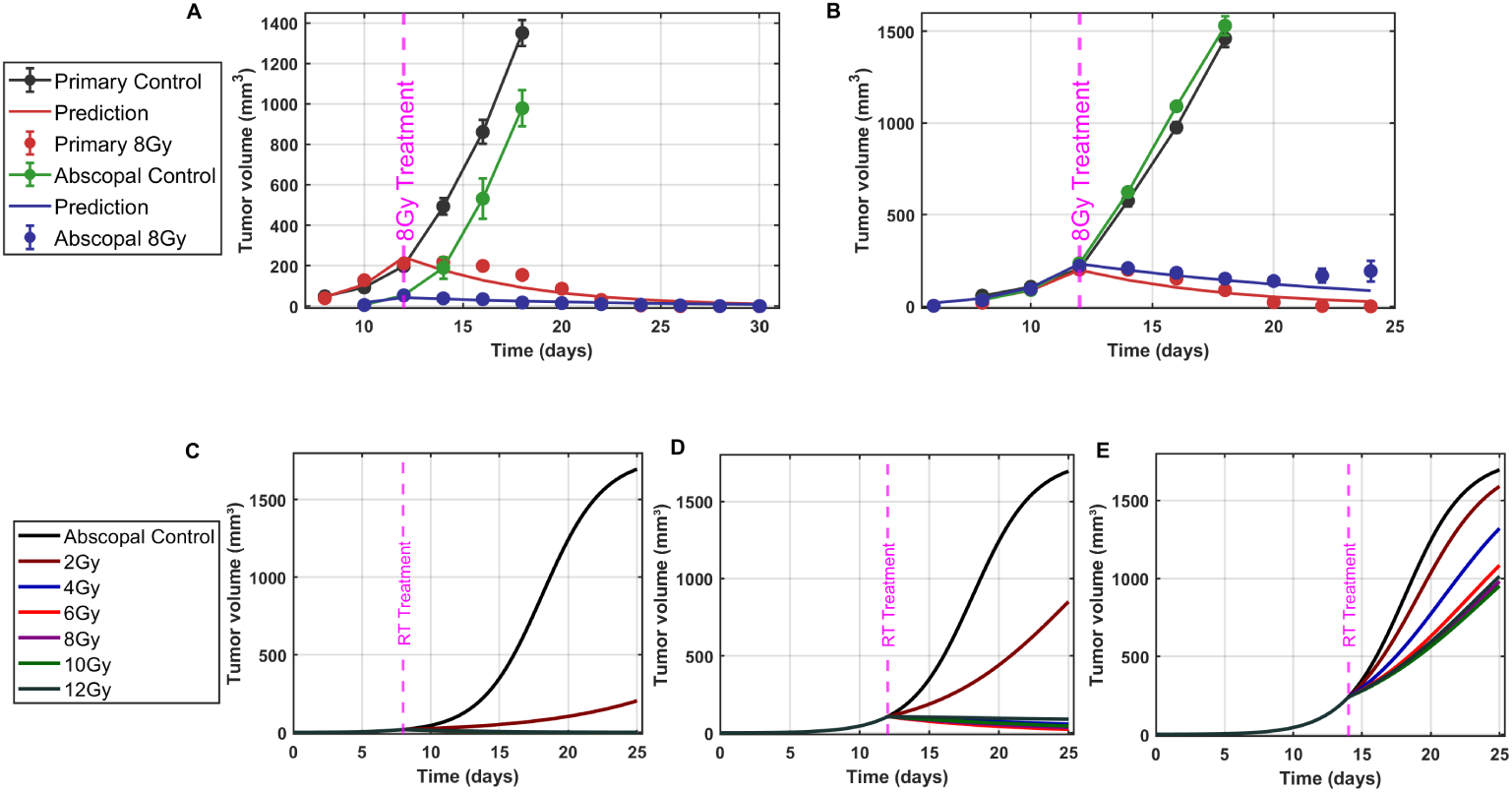
Mathematical model predicts the abscopal effect in MC38 tumors in SIRP*α^−/−^* mice: Our model accurately simulates the abscopal effect in MC38 tumors with different initial engraftments, demonstrating the robustness of its prediction for systemic tumor response to localized radiation therapy. **A.** Initial engraftment of 1 *×* 10^5^ MC38 cells. **B.** Initial engraftment of 5 *×* 10^5^ cells. **C - E.** Predicts abscopal effect for small (C), medium (D) and large (E) tumors across varying RT doses. A medium dose (*≈* 6 Gy) is most effective for reducing small and medium abscopal tumors, while higher doses yield better outcomes for large tumors.

Additionally, the model predicts abscopal effect across different tumor volumes (Figures 7C-E), when the primary tumor is exposed to radiation doses between 2 and 12 Gy. Among these doses, the mid-range dose of *≈* 6 Gy stands out as most effective in reducing the small (*<* 50*mm*^3^, Figure 7C) and medium-sized (50 *−* 200*mm*^3^, Figure 7D) abscopal tumors. For large abscopal tumors (*>* 200*mm*^3^, Figure 7E) higher RT doses (6 Gy) yield better outcomes, mirroring the response observed in primary tumors (Figures 3 and 7E).

#### Treatment efficacy depends on radiation dose and immunogenic activation

We analyze the effects of varying radiation dose (*d*) for different values of the immune activation (*A*) and macrophages phagocytosis (*ϕ_m_*) on the efficacy of tumor treatment. Treatment efficacy is defined as the percent reduction in tumor volume *C*:

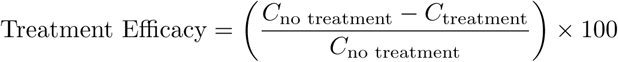

Using the mathematical model, we simulate tumor growth dynamics varying *ϕ_m_*, *A*, and *d* values, and illustrate the isosurfaces of 25%, 50%, 75% and 95% treatment efficacy in a three-dimension (3D) phase space (Figure 8A). In all these cases, the combined RT and checkpoint inhibition is applied on day 12 (intermediate tumor volume), and treatment efficacy is evaluated on day 22. We then map the specific treatments of SIRP*α*-CD47 inhibition for MC38 tumor cells in WT and SIRP*α*-deficient mice according to their phagocytosis values (*ϕ_m_*) as cross-sections through the 3D space. SIRP*α*-KO and anti-SIRP*α* treatments result in the similar *ϕ_m_* values and are therefore grouped together. Sequentially from left to right in the cross-sectional views, we see all the treatment efficacy lines shift upward, suggesting that the irradiation dose required to achieve the same treatment efficacy increases. In other words, for medium sized tumors (100 *−* 400*mm*^3^), given the same irradiation dose, CD47-KO is more effective in reducing tumor than SIRP*α*-KO and anti-SIRP*α*, which in turn is more effective than anti-CD47. The 3D efficacy map for small (*<* 100*mm*^3^) and large (*>* 600*mm*^3^) are only slightly different (Supplementary Figure 5).

**Figure 8.**
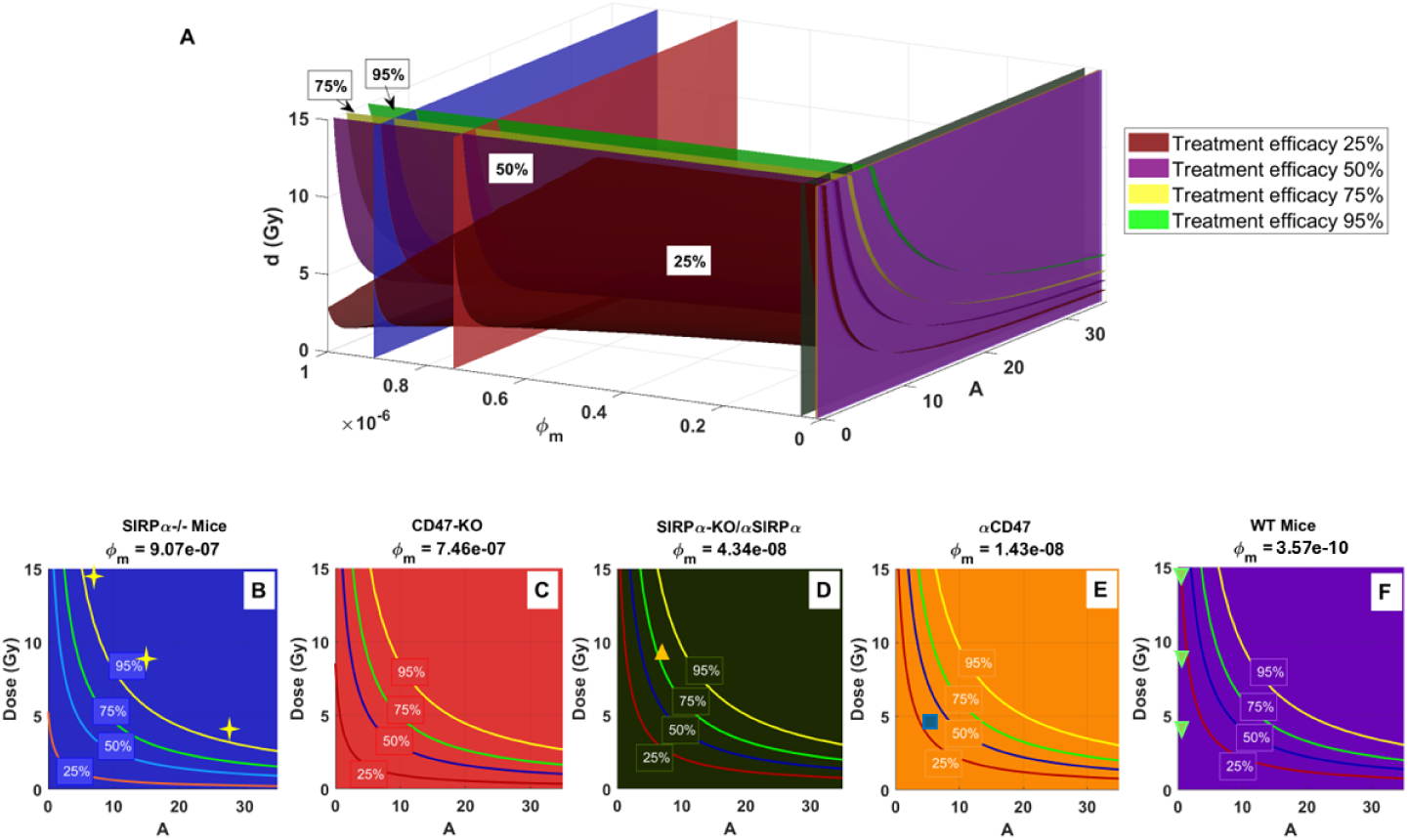
Predicted treatment efficacy for combined RT and macrophage-based immunotherapy: A. Isosurfaces at four efficacy levels (25%, 50%, 75%, and 95%) as a function of ICD activation *A*, macrophage phagocytosis *ϕ_m_*, and radiation dose. B-F: Cross-sections of efficacy contours for *ϕ_m_* values corresponding to various treatment options for inhibiting the SIRP*α*-CD47 checkpoint. Yellow stars in (B) correspond to data in SIRP*α^−/−^* mice with three irradiation doses (data from [25]). Yellow upward triangle in (D) corresponds to twice i.t./i.v. injections of SIRP*α^−^* macrophages into WT mice (data from [25]), blue square in (E) is for anti-CD47 treatment (data from [26]) and green downward triangles in (F) are for WT mice (experimental data from [25]).

A different way to look at the treatment efficacy is presented as a collection of heatmaps for different doses (Supplementary Figure 6. With the model calibrated for MC38 tumors in WT mice, considering both an increased phagocytosis rate *ϕ_m_* due to SIRP*α*-CD47 checkpoint inhibition and an irradiation dose-dependent ICD, we evaluate treatment efficacy when RT is initiated on days 8, 12 and 14, corresponding to small (*<* 100*mm*^3^), intermediate (100 *−* 400*mm*^3^) and large (400 *−* 600*mm*^3^) tumors, respectively. These 2D heatmaps can serve as a guide for optimal RT doses given the tumor size and treatment options.

## Discussion

Cancer cells evade immune destruction through multiple mechanisms, with inhibition of innate recognition and innate-to-adaptive activation of T cell immunity being a major obstacles in immunotherapy. Immunogenic cell death (ICD) has emerged as a key process that enhances antitumor immunity by triggering specific tumor-associated antigens (e.g., CRT, HSP70/90, ATP, and HMGB) [5] and activating immune cells. Our study presents a mathematical model to quantify the role of ICD in enhancing combined RT and macrophage based immunotherapy, specifically targeting the SIRP*α*-CD47 checkpoint. While macrophage based immunotherapies have shown potentials in preclinical murine models, clinical translation remains challenging. CD47 blockade alone has shown tissue toxicity and limited efficacy in human trials [40–42]. SIRP*α* blockade also exhibits unintended phagocytosis of healthy CD47-expressing cells [43, 44], while combination therapies targeting the CD47-SIRP*α* checkpoint show promise [41, 45–47]. These findings underscore the need for optimized combination therapies to enhance efficacy while minimizing safety risks.

Our model, developed through a robust 4-step process, provides several novel insights into the interplay between RT, immune activation, and phagocytosis activation. We find that RT alone fails to generate a sufficient ICD response in WT mice, even at higher doses, highlighting a key limitation of RT-based immunotherapy: without additional immune activation, RT-induced tumor cell death remains largely non-immunogenic. Our model predicts that disrupting the SIRP*α*-CD47 checkpoint enhances macrophage-mediated phagocytosis, shifting the tumor microenvironment toward a more immunostimulatory state. Among different interventions, SIRP*α*-KO macrophages exhibit the highest tumoricidal activity, followed by anti-SIRP*α* and anti-CD47 treatments, suggesting that targeting SIRP*α* directly may offer a more effective strategy than CD47 blockade alone.

Our results identify an optimal radiation dose range (6–8 Gy) that maximizes ICD induction without excessive immune suppression. This aligns with previous experimental findings suggesting that intermediate RT doses enhance tumor immunogenicity, whereas higher doses may impair immune activation. These findings could inform clinical trial design by optimizing radiation schedules to synergize with macrophage-based immunotherapy.

We predict that local RT combined with macrophage activation can induce a systemic immune response, reducing tumor burden beyond the irradiated site (abscopal effect). This result highlights that immune checkpoint inhibitors targeting SIRP*α*-CD47 amplifies the systemic impact of radiotherapy, providing a rationale for combination strategies in metastatic cancer treatment. In these combined RT and SIRP*α*-CD47 checkpoint inhibition treatments, through quantification of enhanced immunogenic activities and ICD, we demonstrate that the primary role of macrophages is the induction of a systemic immune response and the promotion of T cell cytotoxicity that produces additional ICD, not direct tumor killing. Notably, this framework provides a predictive tool for optimizing combination therapies, refining radiation dosing, and quantifying systemic immune responses.

Despite these advances, several challenges remain. While our model captures key aspects of tumor-immune dynamics, further refinements incorporating adaptive immune responses (e.g., T-cell priming and antigen presentation dynamics) would enhance predictive power. Additionally, translating these findings to human tumors requires further validation, as tumor heterogeneity and immune landscape variations may influence ICD induction and treatment efficacy.

Our model parameters align with those reported in tumor dynamics studies. In addition to quantifying the effects of radiation and immunotherapy, our model critically evaluates the role of macrophages type in tumor dynamics. Unlike mathematical models that classify macrophages into distinct M1 and M2 subtypes, our approach accounts for the dynamically altered phagocytosis rates resulting from the perturbations in the CD47-SIRP*α* checkpoint. This perspective aligns with recent findings that view macrophage phenotypes as a continuous and context-dependent spectrum rather than a rigid, bipolar M1/M2 classification [48–50].

To improve the translational relevance of our findings, future studies can focus on experimental validations in human tumor models by expanding preclinical datasets to patient-derived xenografts and organoid models, and assessing how tumor-intrinsic factors affect ICD response. Another important area of future model development is to incorporate adaptive immunity, particularly immune memory formation, to study longer term cancer immunity as reported in [25, 27]. In addition, we can investigate how ICD and macrophage activation interacts with checkpoint inhibitors targeting the PD-1/PD-L1 axis.

This study provides a quantitative framework for understanding and optimizing ICD-driven cancer therapy. By demonstrating that SIRP*α*-CD47 checkpoint inhibition enhances RT-induced ICD, we highlight a potential strategy for improving combination immunotherapies. These findings lay the groundwork for rational treatment design and provide insights into the systemic immune effects of radiotherapy.

## Methods

### Mathematical model of *in vivo* tumor growth with combination therapy

Given the extensive availability of *in vivo* tumor growth data with combination immuno- and radio-therapy [25], we aim to develop a mathematical model that can reproduce and explain these data and predict new treatments. The model construction follows a four-step process. 1. Baseline tumor growth in WT mice: develop a model for *in vivo* tumor growth in wild type mice, calibrating parameters using the experimental data. 2. Tumor growth in SIRP*α*^−*/*−^ mice: extend the baseline model by incorporate SIRP*α*^−^ macrophages to simulate tumor growth in knockout mice. 3. Radiotherapy in WT mice: integrate radiation-induced tumor cell death into the baseline WT tumor model. 4. Radiotherapy in SIRP*α*^−*/*−^ mice: combine the models from steps 2 and 3, incorporating ICD activated by SIRP*α*^−^ macrophages.

The model assumes that radiation-damaged tumor cells release DAMPs, which activate SIRP*α*-deficient macrophages, triggering a systemic pro-inflammatory response that enhances tumor clearance.

#### Step 1: Tumor growth in WT mice

Various models can describe *in vivo* tumor growth, ranging from simple logistic growth models to complex immune-interaction models. We choose the simplest model that best fit the data (details in Supplementary Materials and Supplementary Table 2).

The tumor-immune interaction is modeled as:

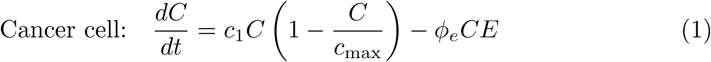

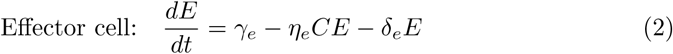

The cancer cell population *C* grows logistically at a rate *c*_1_ with carrying capacity *c*_max_, while *ϕ_e_* represents cancer cell clearance by effector cells *E*. The tumor associated effector cells *E* infiltrate the tumor at a rate rate *γ_e_*, and are cleared at an intrinsic rate of *δ_e_* and an exhaustion rate *η_e_* from combating the tumor cells.

Tumor and effector cell parameters are fitted using WT tumor growth data and kept constant in subsequent steps.

#### Step 2: Modeling tumor growth in SIRP***α*^−^*^/^*^−^** and CD47-SIRP***α*** treated **mice**

To model tumor dynamics in SIRP*α*^−*/*−^ or CD47/SIRP*α*-targeted treatments, we introduce a new cell population *M* ^∗^ for treated macrophages, which exhibit enhanced phagocytosis.

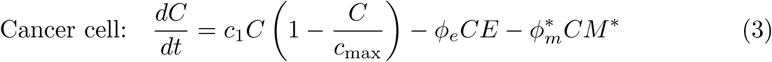

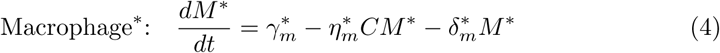

where *ϕ*^∗^_*m*_ is phagocytosis rate. Similar to the effector cells, macrophages with CD47-SIRP*α* treatment has a tumor infiltration rate *γ*^∗^_*m*_, an exhaustion rate *η*^∗^_*m*_, and a natural decay rate *δ*^∗^_*m*_. The effector cell dynamics remains the same as Equation (2).

Four additional parameters related to SIRP*α*-deficient macrophage dynamics fitted using SIRP*α*^−*/*−^ and CD47-SIRP*α* treated tumor growth data.

#### Step 3: RT in WT mice

The impact of RT administered at *t*_1_ is modeled by modifying the tumor cell growth as:

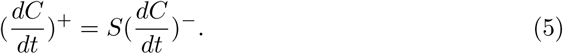

The survival fraction *S* follows the linear-quadratic (LQ) model as a function of radiation dose *d* [36]:

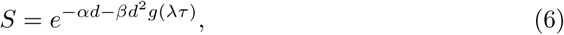

where *α* and *β* are the susceptibility to direct DNA damage and complex damage dynamics involving repair processes, respectively, and *g*(*λτ*) = 2(*λτ* + *e*^−*λτ*^ *−* 1)*/*(*λτ*)^2^ is the repair function, *τ* is the RT delivery time (an experimental constant), and *λ* is the repair parameter [36]. LQ model parameters are fitted using WT tumor growth data under RT.

#### Step 4: RT in SIRP***α*^−^*^/^*^−^** mice or CD47/SIRP***α*** treated mice

In SIRP*α*^−*/*−^ mice or CD47/SIRP*α*-treated mice, increased tumor cell death is attributed to ICD, modeled as:

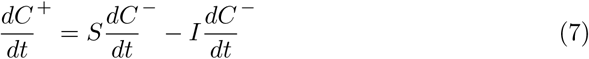

where *I* = *A*(1 *−S*) represents immunogenic cell death and *A* is the immune activation rate. All the other equations and their parameters remain the same as in Steps 2 and 3. The only fitting parameter is the immune activation parameter A for different RT doses and tumor volumes.

Detailed descriptions of model development and parameter estimation are in Supplementary Information. All parameters fall within ranges reported in the literature (Supplementary Table 3).

### Parameters uncertainty quantification

To assess parameter uncertainty, we employ a parametric bootstrapping strategy, assuming that the time-series data follows a Poisson distribution centered on model predictions at each time point [51]. We generate bootstrap samples by randomly sampling *n* = 2000 times (with replacement) from the original data points (as outlined in [52]). The model is then fitted to the bootstrapped datasets using the least squares method, yielding 2000 sets of parameter estimates. From this distribution, we compute the mean and associated uncertainty for each parameter.

### Sensitivity analysis

We perform a local sensitivity analysis to assess how small variations in each parameter affect the model’s output. Parameter ranges are determined through uncertainty quantification, their mean values are the best-fit estimates. To assess sensitivity, each parameter is independently perturb by 25% of its best-fit value, and the resulting relative change in the cost function is measured. The average sensitivity of each parameter is computed across all data points, which forms the basis for a sensitivity ranking that identifies the most and least influential parameters.

## Acknowledgments

We thank Dr. Heiko Enderling for useful discussions on modeling radiotherapy and Dr. Nick Cogan on sensitivity analysis. We appreciate the MBD PhD Fellowship from Georgia State University for S.R., NSF awards 2011622 and 2409868 (DMS) for A.S. and the Frady Whipple Endowed Professorship for Y.J.. This work was partly supported by the institutional funds from the Winship Cancer Institute of Emory University, the NIH 1K08AI178093, and the K12CA237806 from the Emory K12 Clinical Oncology Training Program. Additional support was provided by the Georgia Clinical & Translational Science Alliance under NIH Award UL1TR002378. The content is solely the responsibility of the authors and does not necessarily represent the official views of the National Institutes of Health or National Science Foundation.

