## Supplementary information for "Immunogenic Cell Death: the Key to Unlocking the Potential for Combined Radiation and Immunotherapy"

#### Model development

"Everything should be made as simple as possible, but not simpler." – Albert Einstein

##### Tumor growth in WT mouse: evidence of limited macrophage phagocytosis

We strive to develop a simplest mathematical model that can explain the complex dynamics of *in vivo* tumor growth with immunotherapy and radiotherapy (RT). In our step-by-step model development, we start with a basic ODE model that captures the dynamic interactions between immune and tumor cells, adhering to the methodologies and principles established in prior studies [1–3]. Our first step is to accurately describe *in vivo* tumor growth observed in WT mice, with the influence of immune cells within the tumor microenvironment (TME). When we consider the WT macrophages $M$ as well as the effector cells $E$, we have

$$\begin{aligned}
&\text{Cancer:} && \frac{dC}{dt} = c_1 C\left(1 - \frac{C}{c_{\max}}\right) - \phi_e CE - \phi_{m'} CM,\\
&\text{Effector:} && \frac{dE}{dt} = \gamma_e - \eta_e CE - \delta_e E, \\
&\text{WT Macrophage:} && \frac{dM}{dt} = \gamma_m - \eta_m CM - \delta_m M.
\end{aligned} \tag{S 1}$$

The cancer cell population follows logistic growth with a rate $c_1$ and a carrying capacity $c_{\max}$, while tumor clearance is mediated by effector cells $E$ and WT macrophages $M$, with respective killing rates $\phi_e$ and $\phi_{m'}$. Effector cells and WT macrophages have similar dynamics, characterized by tumor infiltration rates $(\gamma_e, \gamma_m)$, exhaustion rates $(\eta_e, \eta_m)$, and natural clearance rates $(\delta_e, \delta_m)$. In the experiment, varying numbers of MC38 cells ($5 \times 10^3$, $5 \times 10^4$, $5 \times 10^5$, or $2 \times 10^6$) were injected subcutaneously (s.c.) into WT and SIRP$\alpha^{-/-}$ mice [4]. The majority of the injected tumor cells die within 24-48 hours due to the host immune response and the challenging new microenvironment, leaving only a subset that survives and successfully establishes tumor growth [5]. The initial number of engrafted cancer cells after s.c. injection is critical for subsequent tumor growth. To accurately represent the growth dynamics across these four tumor sizes, in addition to the parameters in the equation set S 1, we also fit for the initial tumor size and the initial effector cells and macrophage

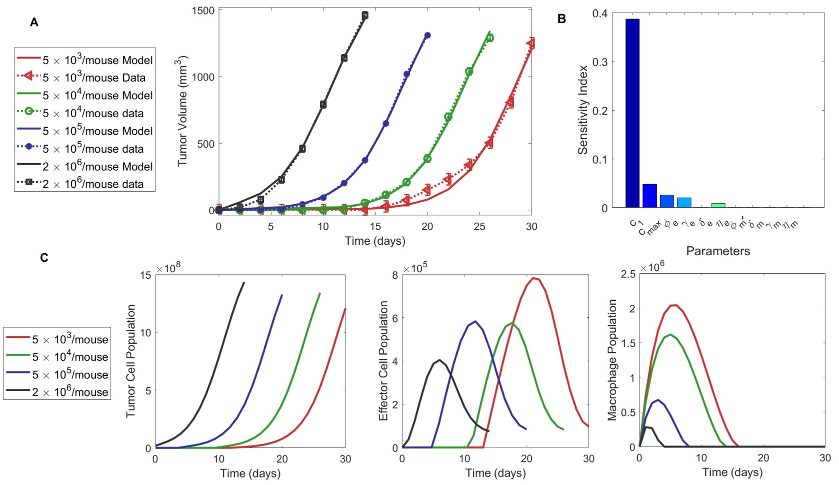


**Supplementary Figure 1** **A.** Fitting the 3D tumor growth model to experimental data, with colors indicating different subcutaneous (s.c.) MC38 injection numbers in WT mice. Dotted line with markers represent experimental data, while solid lines show model fits. **B.** Sensitivity analysis demonstrating the minimal impact of macrophage-specific parameters, suggesting they may be non-essential. **C.** Dynamics of tumor cells, effector cells and macrophages (left to right), with colors corresponding to MC38 cell injection numbers in WT mice.

populations in the tumor. Tumor volume comprises cancer cells, effector cells, and macrophages.

We begin the parameter estimation process by generating 2000 bootstrap samples through resampling from the original data points with replacement. We then fit the model to each bootstrapped dataset using the least squares method (`lsqcurvefit` function in MATLAB, with the cost function is defined as the sum of error squared). This fit produces 2000 sets of parameter estimates. For each parameter, we take the mean of the 2000 estimates as the best-fit value and determine the 95% confidence intervals from the distribution of estimates, summarized in Supplementary Table 1. The model's goodness of fit is evaluated by the error rate defined as |fit - data|/data. This three-compartment (3D) model, which includes WT macrophage, achieves a fit comparable to the 2D model that only considers the tumor cells and effector cells (Supplementary Figure 1A), with error rates of 5.2% and 4.3%, respectively. However, the fitted macrophage killing rate is an order of magnitude lower than reported values for macrophage phagocytosis, while the macrophage clearance rate is 20-times higher (Supplementary Table 1), suggesting that WT macrophages exhibit limited phagocytotic capability in the observed tumor growth data. Moreover, the low sensitivity of macrophage-related parameters suggest that they contribute minimally to model dynamics (Supplementary Figure 1B). It aligns with the understanding that tumor associated macrophages (TAMs), despite their volumetric significance, are more M2-like and do not directly engage in cytotoxic activities within the

**Supplementary Table 1** Parameter estimation for *in vivo* tumor growth in WT mouse (3D model with macrophage compartment).

| **Parameter (Symbol)** | **Unit** | **Literature Value** | **Fitted Value** | **95% CI** |
|---|---|---|---|---|
| Tumor growth rate ($c_1$) | $day^{-1}$ | 4.56 $\times 10^{-1}$ | 4.14 $\times 10^{-1}$ | [3.5e-1, 6e-1] |
| Tumor carrying capacity ($c_{max}$) | $cell$ | $2 \times 10^9$ | $1.91 \times 10^9$ | [1.8e9, 2.1e9] |
| Effector cells killing rate ($\phi_e$) | $(cell \cdot day)^{-1}$ | $1.0 \times 10^{-7}$ | $3.21 \times 10^{-7}$ | [1e-8, 1e-6] |
| Effector cell influx rate ($\gamma_e$) | $cell \cdot day^{-1}$ | $1.3 \times 10^4$ | $1.48 \times 10^4$ | [1e3, 2.53e4] |
| Effector cell exhaustion rate ($\eta_e$) | $(cell \cdot day)^{-1}$ | $3.34 \times 10^{-10}$ | $3.31 \times 10^{-10}$ | [1e-10, 1e-9] |
| Effector cell clearance rate ($\delta_e$) | $day^{-1}$ | $2 \times 10^{-2}$ | $5.9 \times 10^{-2}$ | [2e-3, 2e-1] |
| **Macrophage killing rate ($\phi_{m'}$)** | $\boldsymbol{(cell \cdot day)^{-1}}$ | $\mathbf{6 \times 10^{-9}}$ | $\mathbf{3.57 \times 10^{-10}}$ | **[6e-12, 6e-9]** |
| Macrophage influx rate ($\gamma_m$) | $cell \cdot day^{-1}$ | $1 \times 10^{-7}$ | $3.93 \times 10^{-7}$ | [1e-8, 1e-6] |
| Macrophage exhaustion rate ($\eta_m$) | $(cell \cdot day)^{-1}$ | $1 \times 10^{-7}$ | $1.77 \times 10^{-7}$ | [1e-8, 1e-6] |
| Macrophage clearance rate ($\delta_m$) | $day^{-1}$ | $2 \times 10^{-1}$ | $2.17 \times 10^{-2}$ | [2e-3, 8e-1] |

TME [6]. Taken together, we simplify the *in vivo* tumor growth model by excluding WT macrophage-mediated phagocytosis, reducing model complexity without compromising accuracy.

#### Evaluation of tumor growth models

As many mathematical models can describe tumor growth dynamics, we further examine if the logistic growth model provides the best fit to the data.

These models are evaluated for their ability to fit the *in vivo* tumor growth data (Supplementary Table 2). The Gompertz model exhibits the highest error rate (41.4%), indicating its limitations in this context. The Logistic, Blumberg's, Richards', and Generalized Logistic provide moderate fits with comparatively low error rates. The Exponential model shows a relatively higher error rate (11.6%), suggesting that its simplicity may not adequately capture the complex tumor growth dynamics. Models incorporating strong and weak Allee effects offer additional insights into tumor-immune interactions, emphasizing the role of effector cell density in effective tumor suppression. Ultimately, the Logistic growth model is chosen, as it strikes a balance between complexity and accuracy while minimizing the risk of over-fitting.

$$\begin{aligned} \frac{dC}{dt} &= c_1 C \left(1 - \frac{C}{c_{\max}}\right) - \phi_e C E, \\ \frac{dE}{dt} &= \gamma_e - \eta_e C E - \delta_e E. \end{aligned} \tag{S 2}$$

#### Tumor growth in SIRP$\alpha^{-/-}$ mice: enhanced phagocytosis for SIRP$\alpha^{-/-}$ macrophages

To model tumor growth in SIRP$\alpha^{-/-}$ mice, we introduce SIRP$\alpha$-deficient macrophages, denoted as $M^*$. SIRP$\alpha$-deficient macrophages show enhanced tumor cell

**Supplementary Table 2  Comparison of 2D models for tumor growth in WT mice**

| Model | Equation[a] | Error Rate[b] |
|---|---|---|
| **Generalized Logistic** [7] | $\frac{dC}{dt} = c_1 C^a \left(1 - \left(\frac{C}{c_{\max}}\right)^b\right)^c - \phi_e CE$ | 5.0% |
| **Richards'** [7] | $\frac{dC}{dt} = c_1 C \left(1 - \left(\frac{C}{c_{\max}}\right)^b\right) - \phi_e CE$ | 4.7% |
| **Blumberg's** [7] | $\frac{dC}{dt} = c_1 C^a \left(1 - \frac{C}{c_{\max}}\right)^c - \phi_e CE$ | 4.4% |
| **Logistic** [7] | $\frac{dC}{dt} = c_1 C \left(1 - \frac{C}{c_{\max}}\right) - \phi_e CE$ | 4.3% |
| **Exponential** [7] | $\frac{dC}{dt} = c_1 C - \phi_e CE$ | 11.6% |
| **Gompertz** [7] | $\frac{dC}{dt} = c_1 C \log\left(\frac{c_{\max}}{C}\right) - \phi_e CE$ | 41.4% |
| **Strong Allee Effect** [8] | $\frac{dC}{dt} = c_1 C \left(1 - \frac{C}{c_{\max}}\right) - \phi_e CE \left(\frac{E}{A_l} - 1\right)$ | 4.42% |
| **Weak Allee Effect** [8] | $\frac{dC}{dt} = c_1 C \left(1 - \frac{C}{c_{\max}}\right) - \phi_e CE \left(\frac{E}{A_l} - 1\right)$ | 4.8% |

a.The effector equation $\frac{dE}{dt} = \gamma_e - \delta_e E - \eta_e CE$ remains the same for all models. b. Error rate is defined as |fit - data|/data.

**Supplementary Table 3  Parameter estimation for tumor growth in WT mice (2D model): All parameter values fall within ranges of literature-reported values.**

| Parameter (Symbol) | Unit | Literature Value | Fitted Value (Mean) | 95% CI |
|---|---|---|---|---|
| Tumor growth rate ($c_1$) | $day^{-1}$ | $4.56 \times 10^{-1}$ [9] | $4.53 \times 10^{-1}$ | [3.8$e$−1, 6$e$−1] |
| Tumor carrying capacity ($c_{\max}$) | $cell$ | $2 \times 10^9$ [9] | $1.77 \times 10^9$ | [1.7$e$9, 2.1$e$9] |
| Effector cells killing rate ($\phi_e$) | $(cell \cdot day)^{-1}$ | $1.0 \times 10^{-7}$ [10] | $4.5 \times 10^{-8}$ | [2$e$−8, 9.4$e$−8] |
| Effector cell influx rate ($\gamma_e$) | $cell \cdot day^{-1}$ | $1.3 \times 10^4$ [1] | $1.65 \times 10^4$ | [5.0$e$3, 3.37$e$4] |
| Effector cell exhaustion rate ($\eta_e$) | $(cell \cdot day)^{-1}$ | $3.34 \times 10^{-10}$ [10] | $1.98 \times 10^{-9}$ | [3.3$e$−10, 3.3$e$−9] |
| Effector cell clearance rate ($\delta_e$) | $day^{-1}$ | $2 \times 10^{-2}$ [1] | $7.41 \times 10^{-2}$ | [6.8$e$−3, 2$e$−1] |

phagocytosis. The equation for $M^*$ is similar to that of WT macrophage, incorporating a tumor infiltration rate, an exhaustion rate, and a baseline clearance rate:

$$\begin{aligned} \text{Cancer:} \quad & \frac{dC}{dt} = c_1 C \left(1 - \frac{C}{c_{\max}}\right) - \phi_e CE - \phi_m^* CM^*, \\ \text{Effector:} \quad & \frac{dE}{dt} = \gamma_e - \eta_e CE - \delta_e E, \\ \text{SIRP}\alpha^{-/-} \text{ Macrophage:} \quad & \frac{dM^*}{dt} = \gamma_m^* - \eta_m^* CM^* - \delta_m^* M^*. \end{aligned} \tag{S 3}$$

To fit the parameters for tumor growth in SIRP$\alpha^{-/-}$ mice, we keep the six parameters associated with the tumor cells and effector cells unchanged and focus solely on fitting the four parameters related to macrophages using the same uncertainty quantification analysis. We find that SIRP$\alpha^{-/-}$ macrophages exhibit superior phagocytic activity (Supplementary Table 4), two orders magnitude higher than the reported value for M1 macrophages [11]. This observation agrees with experimental findings [4, 12] which report the enhanced phagocytic capability of SIRP$\alpha^{-/-}$ macrophages. This heightened phagocytic capability contributes to the observed deceleration in tumor growth when exposed to a lower inoculum, highlighting the enhanced immune control exhibited by SIRP$\alpha^{-/-}$ mice *in vivo*.

The calibrated model enables us to separately show the contributions of cancer cell, effector cell, and macrophages to the tumor volume (Supplementary Figure 2).

**Supplementary Table 4** Parameters estimation for tumor growth in SIRP$\alpha^{-/-}$ mice with initial conditions $4.89 \times 10^6$, $2.80 \times 10^7$, $2.69 \times 10^7$, $5.77 \times 10^7$, for tumor engraftments (from low to high), and $4.30 \times 10^6$, and $3.43 \times 10^5$ for effector cells and SIRP$\alpha^{-/-}$ macrophages respectively.

| **Parameter (Symbol)** | **Unit** | **Literature Value (M1 macrophage)** | **Fitted value (Mean)** | **95% CI** |
|---|---|---|---|---|
| $M^*$ killing rate ($\phi^*_m$) | $(cell \cdot day)^{-1}$ | $6 \times 10^{-9}$[11] | $9.07 \times 10^{-7}$ | [1e−7, 1e−5] |
| $M^*$ influx rate ($\gamma^*_m$) | $cell \cdot day^{-1}$ | $1 \times 10^{-7}$[9] | $4.39 \times 10^{-7}$ | [1e−8, 1e−6] |
| $M^*$ exhaustion rate ($\eta^*_m$) | $(cell \cdot day)^{-1}$ | $1 \times 10^{-7}$[11] | $8.15 \times 10^{-7}$ | [1e−8, 1e−6] |
| $M^*$ clearance rate ($\delta^*_m$) | $day^{-1}$ | $2 \times 10^{-1}$[9] | $3.94 \times 10^{-1}$ | [2e−3, 8e−1] |

$M^*$ denotes SIRP$\alpha^{-/-}$ macrophage.

#### Modeling RT

For RT applied to tumors in WT mice, we start with the WT tumor growth model S 2 and introduce RT as a stepwise modification to the continuous equation. Instead of assuming instantaneous tumor cell elimination upon application, we account for the prolonged effects of RT, as its impact extends over time [13].

To investigate tumor response across various radiation doses, we incorporate RT-induced cancer cell death using a linear-quadratic dose-dependent survival fraction [14, 15].

$$S = e^{-\alpha d - \beta d^2 g(\lambda\tau)}, \tag{S 4}$$

where $\alpha$ and $\beta$ are the susceptibility to direct DNA damage and complex damage dynamics involving repair processes, respectively, and $g(\lambda\tau)$ is the repair function:

$$g(\lambda\tau) = 2\frac{\lambda\tau + e^{-\lambda\tau} - 1}{(\lambda\tau)^2}, \tag{S 5}$$

where $\tau$ is the duration of radiation delivery and $\lambda$ is the repair parameter [14]. Reported radiation delivery rates vary across studies, such as 1.2 Gy/min in [4],

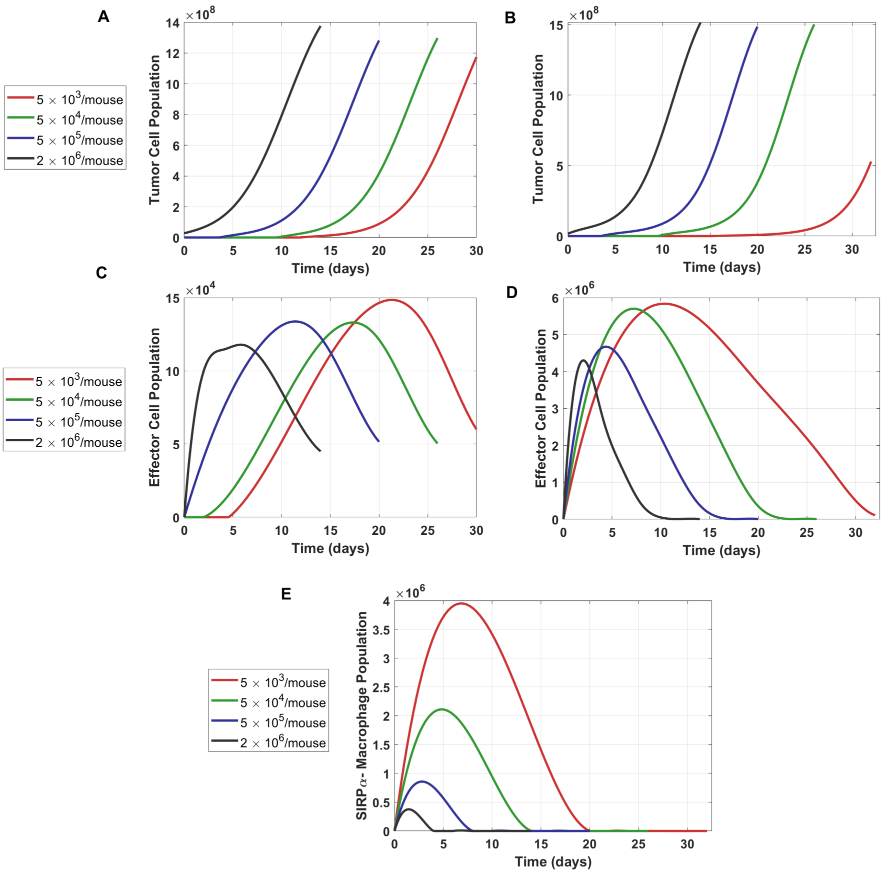


**Supplementary Figure 2 Tumor growth in WT and SIRP$\alpha^{-/-}$ mice. A.** Tumor growth across four engraftment levels of MC38 cells in WT mice. **B.** Tumor growth is slower in SIRP$\alpha^{-/-}$ mice for low engraftment level (red line) but otherwise indistinguishable from WT mice. **C.** Effector cell dynamics in WT mice, showing a peak when tumor volume approaches $200mm^3$ (corresponding part A). **D.** Effector cell dynamics in SIRP$\alpha^{-/-}$ mice, with a higher maximum for low engraftment (red line). **E.** SIRP$\alpha$-deficient macrophages in tumor reach higher levels in lower engraftment levels. Experimental data utilized here is from [4].

2.41 Gy/min in [16], 2.53 Gy/min for [17]. We calculate $\tau$ for each dose in each experiment accordingly.

Irradiation activates the immune system, prompting the release of cytokines, chemokines, and growth factors, which play crucial roles in regulating immune responses, inflammation, cell recruitment, tissue growth, and repair [18]. Assuming the amount of DAMPs generated by radiation is proportional to the fraction of damaged cancer cells $(1-S)$ and immune activation is proportional to DAMPs, we introduce an immunogenic death term that scales with $(1-S)$. The cancer cell population post-RT grows at a fraction $S$ of its pre-RT rate and decays at a rate proportional to $1-S$:

$$\frac{dC^+}{dt} = S\frac{dC^-}{dt} - A(1-S)\frac{dC^-}{dt}, \qquad \text{(S 6)}$$

where $A$ is the immune activation rate and we define $I = A(1 - S)$ immunogenic cell death (ICD).

Supplementary Table 5 summarizes the parameters in the linear-quadratic model. The calibrated parameter values show minimal $A$ and $I$ values, suggesting limited ICD in WT mice after RT.

**Supplementary Table 5 Parameters for Linear Quadratic Radiotherapy Model**

| Parameter (Symbol) | Unit | Literature Value | Mean of Fitted Values | 95% CI |
|---|---|---|---|---|
| Linear damage coefficient ($\alpha$) | $Gy^{-1}$ | $2.8 \times 10^{-8}$ [14] | $5.15 \times 10^{-3}$ | [0, $5.4e{-}3$] |
| Quadratic damage coefficient ($\beta$) | $Gy^{-2}$ | $1.32 \times 10^{-2}$ [14] | $6.58 \times 10^{-3}$ | [0, $2.24e{-}2$] |
| Repair parameter ($\lambda$) | $day^{-1}$ | 2.0358 [14] | 2.56 | [2.54, 5.00] |

We combine the model for WT tumor growth model (S 2) and for SIRP$\alpha^{-/-}$ tumor growth model (S 3) and RT (Supplementary Equations S 4, S 5 and S 6) to model RT in WT and SIRP$\alpha^{-/-}$ mice, respectively. A significantly enhanced ICD is observed for each dose and varying tumor volumes.

### Sensitivity analysis

We use a systematic perturbation approach to assess the sensitivity of parameters, leveraging a least-squares optimization framework for model calibration.

The perturbation factor, denoted by $f$, is set to $f = 0.25$. $x_p$ the original parameter value, and $x_p(p) = x_p \pm \epsilon(p))$ the perturbed value. The sensitivity matrix is $S(p) = \frac{C(x_p(p)) - C(x)}{C(x)}$, where $C(x)$ is the cost value obtained from the mean parameter values computed in uncertainty quantification. The sensitivity index for each parameter is computed as the average of the absolute values in the sensitivity matrix. This index quantifies the relative importance of each parameter in influencing the model predictions.

In the WT model, we find that the tumor cell growth rate, $c_1$, and the maximum tumor capacity, $c_{\max}$, are the most sensitive parameters, affecting tumor growth dynamics significantly, followed by the parameters related to the effective cells $\phi_e$ and $\gamma_e$, while the effector cell exhaustion rate is less sensitive (Supplementary Figure 3B). Parameters associated with macrophages show minimal influence on the model's behavior (Supplementary Figure 1B). On the other hand, in the SIRP$\alpha^{-/-}$ mice, $\eta_m^*$ emerges as a sensitive parameter, underscoring the key role of SIRP$\alpha^{-/-}$ macrophages (Supplementary Figure 3B).

Moreover, when RT is applied, the sensitivity analysis shows the time dependent impact of RT parameters $\alpha, \beta$, and $\lambda$. In WT mice, these parameters exert minimal influence with sensitivity values less than $10^{-3}$ (Supplementary Figure 4A). Conversely, in SIRP$\alpha^{-/-}$ mice, the relative sensitivity RT parameters change over time

**Supplementary Table 6** **Comparison of parameter values across cell lines and mice models**

| **Cell Line/Model [data source]** | $c_1$ ($\times 10^{-1}$) | $c_{\max}$ ($\times 10^{9}$) | $\alpha$ ($\times 10^{-3}$) | $\beta$ ($\times 10^{-3}$) |
|---|---|---|---|---|
| MC38 in WT [4] | 4.53 | 1.77 | 5.15 | 6.58 |
| Pan02 in WT [4] | 4.40 | 2.00 | 5.15 | 6.58 |
| KPC in WT [4] | 4.80 | 2.00 | 5.15 | 6.58 |
| Fibrosarcoma in WT BALB/c [17] | 3.59 | 4.50 | 5.15 | 6.58 |
| NSG KP1 SCLC [16] | 5.10 | 4.08 | $3.80 \times 10^{-6}$ | 25.8 |
| MC38-OVA in WT [19] | 4.53 | 1.77 | 1.00 | 65.0 |
| MC38 in WT [20] | 1.60 | 4.32 | 5.15 | 6.58 |

(Supplementary Figure 4B), highlighting the altered dynamics due to the immune profile modification.

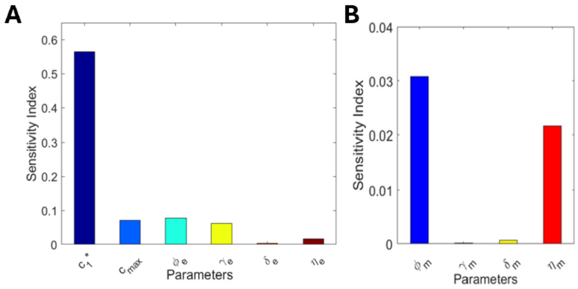


**Supplementary Figure 3** **A.** Sensitivity analysis in WT mice, highlighting the sensitivity of $c_1$, $c_{\max}$, $\phi_e$, and $\gamma_e$. **B.** shows the key role of $\phi_m^*$, $\eta_m^*$ in tumor growth in SIRP$\alpha^{-/-}$ mice. Although the sensitivity of $\eta_m^*$, $\gamma_m^*$ is of negligible order, we chose to retain them in the model because of availability of sufficient data points and our intention to preserve the biological relevance of the system.

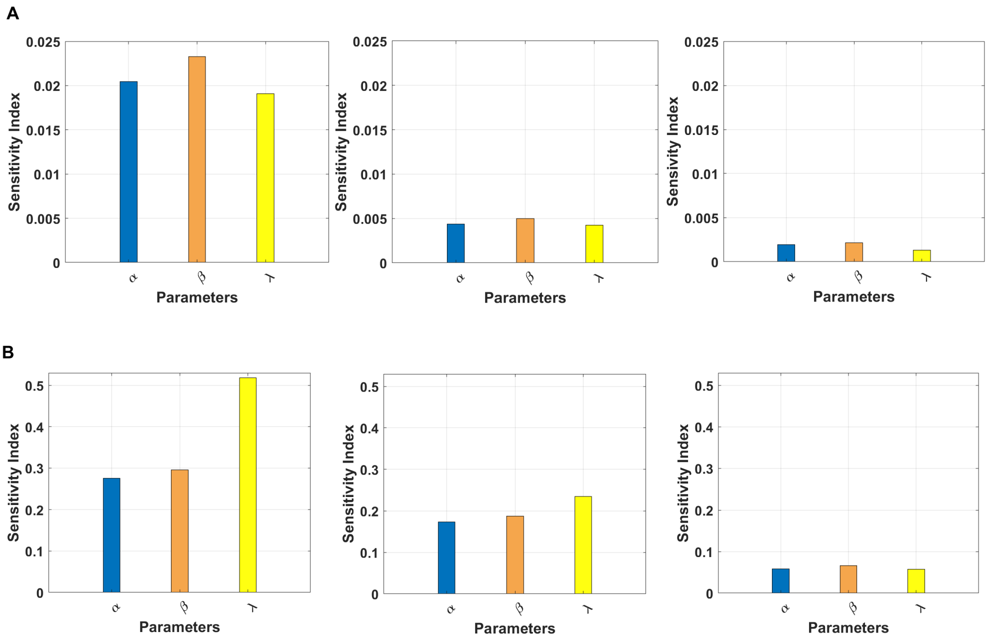


**Supplementary Figure 4** Sensitivity analysis of RT parameters in WT and SIRP$\alpha^{-/-}$ mice across different tumor sizes: **A.** Sensitivity analysis of RT parameters ($\alpha$, $\beta$, and $\lambda$ ) in WT mice for small (left), medium (center), and large (right) tumors. The sensitivity values are in the order of $10^{-3} - 10^{-2}$, indicate minimal influence. **B.** Sensitivity analysis of RT parameters ($\alpha$, $\beta$, and $\lambda$ ) in SIRP$\alpha^{-/-}$ mice, for small (left), medium (center), and large (right) tumors. The relative sensitivity of RT parameters change over time.

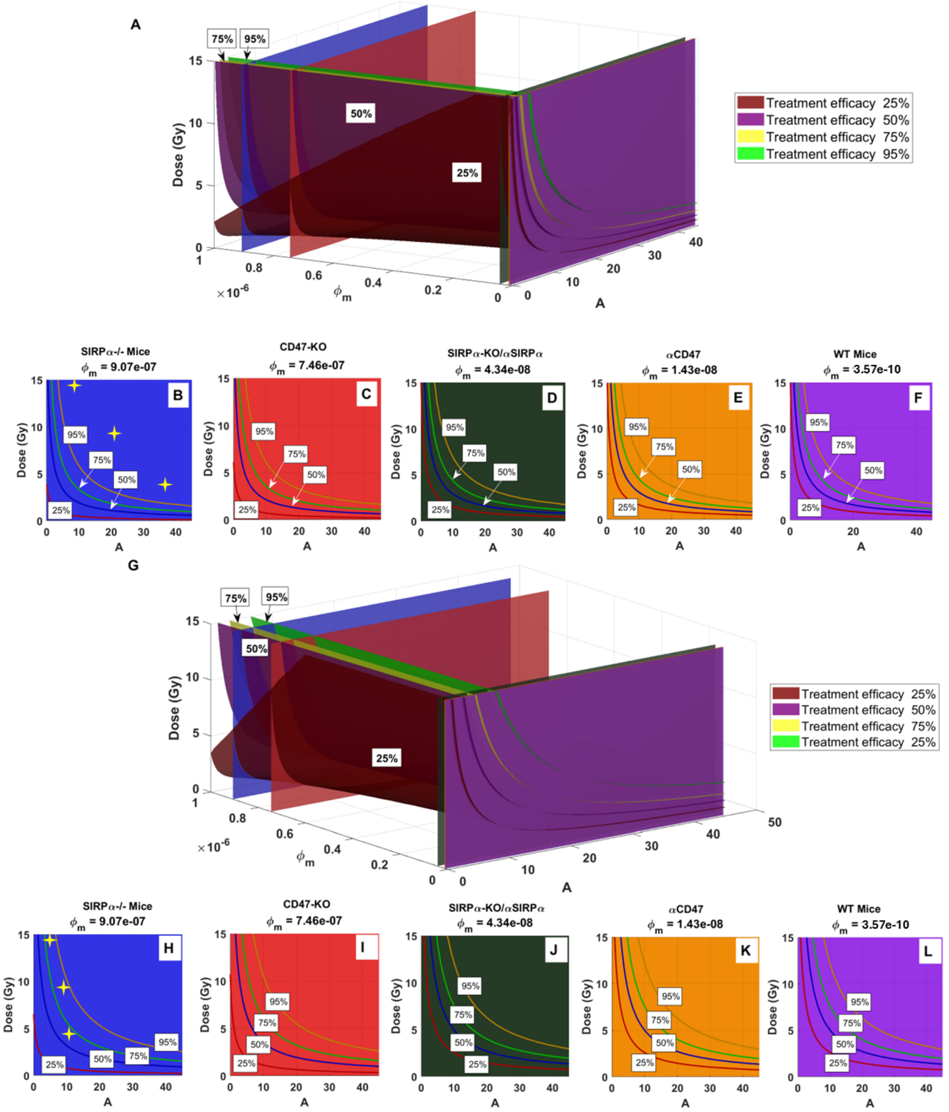


**Supplementary Figure 5 Predicted treatment efficacy for combined RT and macrophage-based immunotherapy for small tumors (A-F) and large tumors (G-L):** A &G. Isosurfaces at four efficacy levels (25%, 50%, 75%, and 95%) as a function of ICD activation $A$, macrophage phagocytosis $\phi_m$, and radiation dose. B-F & G-L: Cross-sections of efficacy contours for $\phi_m$ values corresponding to various treatment options for inhibiting the SIRP$\alpha$-CD47 checkpoint. Yellow stars in (B & H) correspond to data in SIRP$\alpha$/ mice with three irradiation doses (data from [25]).

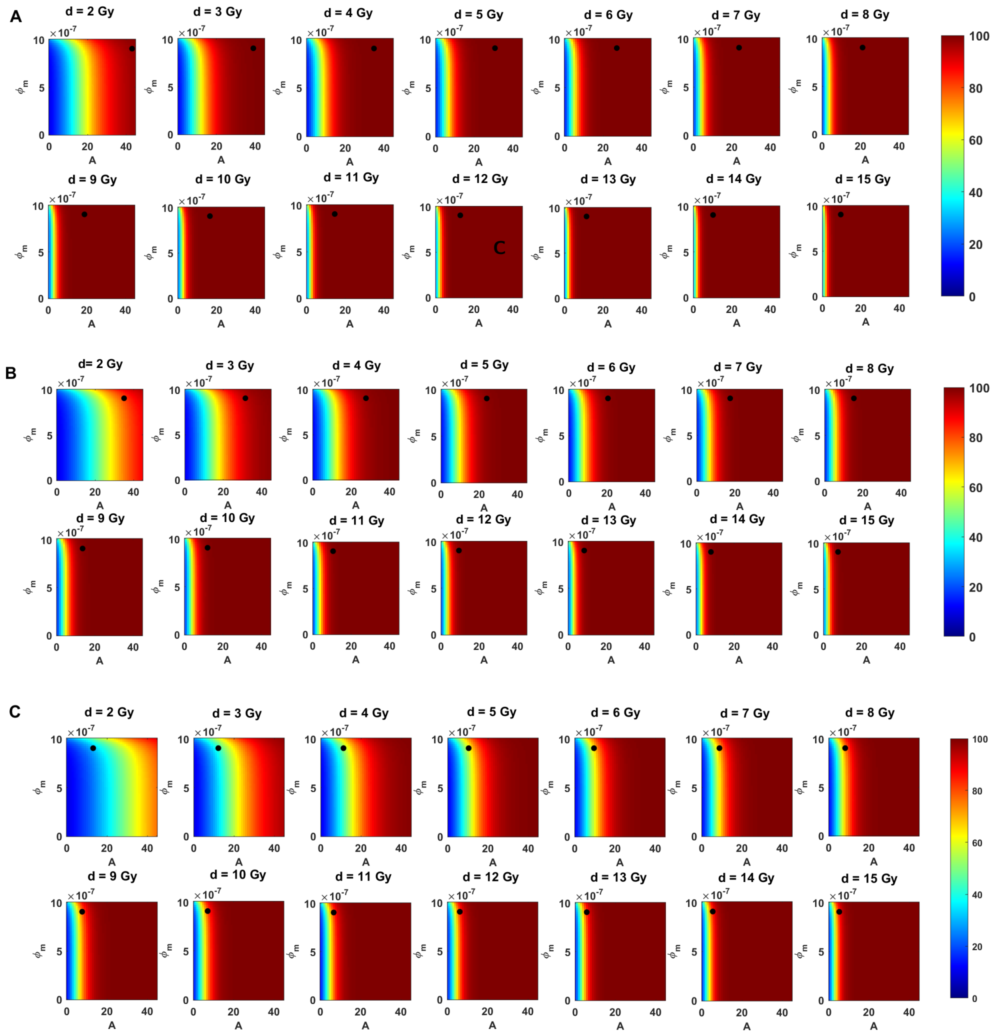


**Supplementary Figure 6 Tumor size dependent response to RT:** The heat maps illustrate the predicted impact of radiation dose, phagocytosis and immune activation on tumor reduction efficacy. (A) Small tumors exhibit high sensitivity to RT, showing significant reduction across all dose levels. (B) Medium-sized tumors demonstrate reduced efficacy at lower doses but retain responsiveness at higher doses. (C) Large tumors display minimal response, particularly at low doses, highlighting the critical role of tumor size and the need for timely intervention to achieve optimal therapeutic outcomes.

**Supplementary Table 7 Overview of experimental setups and modeling parameters across different treatments**

| Mice Model | Cell Line | Description | Data Points | Fitted Parameters | Remarks |
|---|---|---|---|---|---|
| C57BL/6 (WT) | MC38 | Mice were s.c. engrafted with four various initial injections | 32 | $c_1 = 4.53 \times 10^{-1}$, $c_{\max} = 1.77 \times 10^{9}$, $\phi_e = 4.5 \times 10^{-8}$, $\gamma_e = 1.65 \times 10^{4}$, $\eta_e = 1.98 \times 10^{-9}$, $\delta_e = 7.41 \times 10^{-2}$ | Parameters remain within the same orders of magnitude reported in literature; 7% error. |
| SIRP$\alpha$-/- | MC38 | Mice were s.c. engrafted with four various injections | 32 | $\phi^*_m = 9.07 \times 10^{-7}$, $\gamma^*_m = 4.39 \times 10^{-7}$, $\eta^*_m = 8.15 \times 10^{-7}$, $\delta^*_m = 3.94 \times 10^{-1}$ | Error is 7%; the parameters fitted in WT mice remain constant. |
| C57BL/6 (WT) with RT | MC38 | Mice were s.c. engrafted with $5 \times 10^5$/mouse of MC38 cells, with control, 4, 8, and 15 Gy RT applied to small medium and large tumor | Dataset 1: 35; Dataset 2: 30; Dataset 3: 29 | $\alpha = 5.15 \times 10^{-3}$, $\beta = 6.58\times10^{-3}$, $\lambda = 2.56$ | The parameters fitted in WT mice remain constant. The error rates are 8%, 5%, and 8%. |
| SIRP$\alpha$-/- with RT | MC38 | Mice were s.c. engrafted with $5 \times 10^5$/mouse of MC38 cells, and no RT, 4, 8, and 15 Gy radiation treatments were applied on days 8, 12, and 14 | Dataset 1: 37; Dataset 2: 37; Dataset 3: 34 | $A = 34.9, 21.1, 9.6$; $A = 27.7, 15.3, 7.7$; $A = 11.3, 8.2, 5.3$ | The error rates are 5%, 8%, and 6%. The parameters fitted in WT mice and WT RT remain constant. |
| SIRP$\alpha$-/- with RT | MC38 | Disrupting SIRP$\alpha$-deficient macrophages using Cl2MDA liposomes or an antibody against the CSF1 receptor ($\alpha$CSF1R) from SIRP$\alpha$-/- mice, then 8 Gy RT was applied on day 12 | 36 | All previous values used | All other parameters remain constant. |
| WT with RT | MC38 | The i.v. injection of SIRP$\alpha$-deficient macrophages into WT, then 8 Gy RT was applied on days 12 and 14 | 18 | $A_1 = 8.247$, $A_2 = 19.95$ and $\phi^*_m = 4.43 \times 10^{-8}$ | All other parameters remain constant. |
| WT with RT | MC38 | The i.t. injection of BMDM and SIRP$\alpha$-/- macrophages injection into WT; 8 Gy RT applied on day 12 | 34 | $A$ values from i.v. case is used | 7.2% error rate is observed. All parameters remain constant. |
| WT with RT | MC38 | The i.t. injection of BMDM and SIRP$\alpha^{-/-}$ macrophages into WT; 8 Gy RT applied on days 12 and 14 | 34 | $A$ values from i.v. case is used | 9.9% error rate is observed. All parameters remain constant. |
| WT+SIRP$\alpha$-/- with RT | KPC | (s.c.) injected KPC into WT and SIRP$\alpha$-/-; 8 Gy RT applied on day 18 | 31 | $c_1 = 4.8 \times 10^{-1}$, $c_{max} = 2\times10^{9}$, $A = 17.24$ | 4.9% error rate. All other parameters remain constant. |
| WT+SIRP$\alpha$-/- with RT | Pan02 | (s.c.) injected Pan02 into WT and SIRP$\alpha$-/-; 8 Gy RT applied on day 12 | 32 | $c_1 = 4.4 \times 10^{-1}$, $c_{max} = 2\times10^{9}$, $A = 15.33$ | 3.9% error rate is observed. All other parameters remain constant. |
| WT BALB/c Mice with RT | 15-12RM fibrosarcoma | Intraperitoneally (i.p.) injected into Athymic BALB/c nu/nu mice with saline as a control or with a solution of 10 mmol/L CD47 morpholino (CD47M) in saline with 10 Gy RT applied on day 10 | 27 | $c_1 = 3.59 \times 10^{-1}$, $c_{max} = 4.5 \times 10^{9}$, $A = 0$ | 8.3% error rate is observed. All other parameters remain constant. |
| Immunodeficient NSG Mice with RT | KP1 SCLC | Anti-CD47 applied. Two datasets with 5 Gy dose applied on days 10 or 12 | Dataset 1: 20; Dataset 2: 20 | $c_1 = 5.1 \times 10^{-1}$, $c_{max} = 4.08 \times 10^{9}$, $\alpha = 3.8 \times 10^{-9}$, $\beta = 2.58 \times 10^{-2}$, $\phi^*_m = 1.43 \times 10^{-8}$, $A = 1.42$, $A = 5.41$ | 4.3% and 11.8% error rate are observed. All other parameters remain constant. |
| Immunodeficient NSG Mice with RT | KP1 SCLC | CD47-KO for two datasets of RT 5 Gy double doses applied on days 10, 14 and 11, 13 | Dataset 1: 22; Dataset 2: 21 | $A_1 = 0.16$, $A_2 = 2.47$; $A_1 = 0.97$, $A_2 = 2.97$ | 14.6% and 15.2% error rates. All other parameters remain constant on the basis of anti-CD47 fitting. |
| WT Mice with RT | MC38-OVA | anti-CD47/anti-SIRP$\alpha$ treatment with 8 Gy applied on days 12 | 32 | $\alpha = 1 \times 10^{-3}$, $\beta = 6.5 \times 10^{-2}$, $\phi^*_m = 4.34 \times 10^{-8}$, $A_{cd} = 0.27$, and $A_{sa} = 0.33$ | 10.6% error, RT delivery time not given in article, we assumed 6Gy/min. All other parameters remain constant on the basis of anti-CD47 fitting. |
| WT Mice with RT | MC38 | Anti-SIRP$\alpha$ treatment with 12 Gy RT applied on day 9 | 32 | $c_1 = 1.6 \times 10^{-1}$, $c_{max} = 4.32 \times 10^{9}$ and $A = 0.99$ | 9.3% error, RT delivery time not given in article, we assumed 6Gy/min. All other parameters remain constant on the basis of anti-SIRP$\alpha$ fitting in WT mice. The reason for fitting $c_1$ and $c_{max}$ is the reduce tumor growth in this study. |
